# On the patterns of genetic intra-tumour heterogeneity before and after treatment

**DOI:** 10.1101/2024.10.10.617575

**Authors:** Alexander Stein, Benjamin Werner

## Abstract

Genetic intra-tumour heterogeneity (gITH) is a universal property of all cancers. It emerges from the interplay of cell division, mutation accumulation and selection with important implications for the evolution of treatment resistance. Theoretical and data-driven approaches extensively studied gITH in ageing somatic tissues or cancers at detection. Yet, the expected patterns of gITH during and after treatment are less well understood. Here, we use stochastic birth-death processes to investigate the expected patterns of gITH across different treatment scenarios. We consider homogeneous treatment response with shrinking, growing and stable disease, and follow up investigating heterogeneous treatment response with sensitive and resistant cell types. We derive analytic expressions for the site frequency spectrum, the total mutational burden and the single-cell mutational burden distribution that we validate with computer simulations. We find that the SFS after homogeneous treatment response retains its characteristic power-law tail, while emergent resistant clones cause peaks corresponding to their sizes. The frequency of the largest resistant clone is subdominant and independent of the population size at detection, whereas the relative total number of resistant cells increases with detection size. Furthermore, the growth dynamics under treatment determine whether the total mutational burden is dominated by preexisting or newly acquired mutations, suggesting different possible treatment strategies.

## Introduction

Every cell in a human body accumulates thousands of unique mutations with age [1, 2]. From this genetic heterogeneity, pre-malignant cells may arise and eventually evolve into an expanding cancerous cell population. As the disease progresses, cancer cells continue to accumulate mutations, subject to genetic drift and selection, leading to a complex pattern of genetic intra-tumor heterogeneity (gITH) [3, 4].

Mathematical and computational modelling applied to sequencing data of normal and cancerous tissues has led to a better understanding of the emerging patterns of clonal diversity [5, 6, 7, 8, 9]. It is clear that gITH informs on past evolutionary paths and at least partially determines future evolution, e.g. the risk of resistance or relapse in treated tumours [10, 11]. While patterns of gITH in both normal and cancerous cell populations prior to treatment are well studied, less is known about the dynamics of gITH in response to treatment.

In this study, we use birth-death processes with random accumulation of mutations to investigate gITH in cancers before and after treatment. We investigate three different measures of gITH: the site frequency spectrum (SFS), the total mutational burden (tMB) and the single cell mutational burden (scMB). We first derive analytic expressions for populations at detection and homogeneous treatment responses. We then investigate effects of emerging resistance and the corresponding heterogeneous treatment response on possible measures of gITH and their dependence on the detailed dynamics and timing of treatment.

## Theoretical framework

We model the cancer cell population using a birth-death process with constant birth and death rates. Starting from a single ancestor cell, cells either divide or die giving rise to a phylogeny (fig. 1A). Initially, the birth rate is larger than the death rate (*b*_1_ > *d*_1_) leading to a growing population (phase 1). Once the cancer is detected at time *t*_*d*_ with size *N*_*d*_, treatment is administered leading to new birth and death rates *b*_2_ and *d*_2_ (phase 2). Treatment is applied for time *t*_*f*_ after which we have the final population size *N*_*f*_. Assuming that all cells have the same birth and death rates, there are three qualitatively different scenarios for the growth during treatment: The cancer decreases (*b*_2_ < *d*_2_ and *N*_*d*_ > *N*_*f*_), remains approximately constant (*b*_2_ = *d*_2_ and *N*_*d*_ ≈ *N*_*f*_) or continues growing (*b*_2_ > *d*_2_ and *N*_*d*_ < *N*_*f*_) as illustrated in fig. 1B.

**Figure 1:**
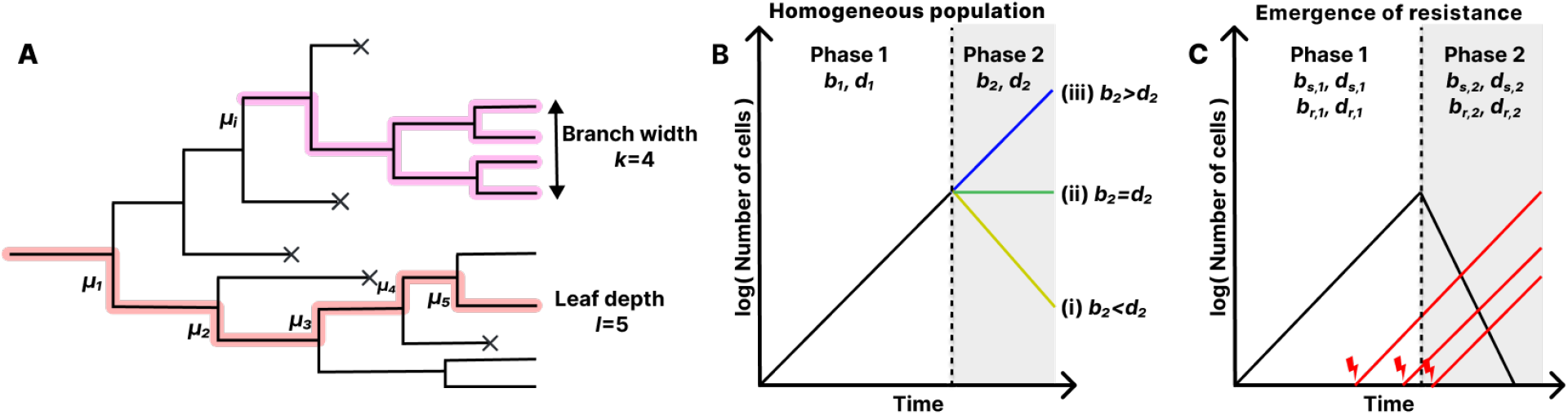
Model illustration. **A.** Phylogeny with highlighted branch (purple) and highlighted path from leaf to root (red). **B**. Growth of homogeneous population: In phase 1, the population grows exponentially. In phase 2, the population (i) decreases, (ii) remains constant or (iii) continues increasing. **C**. Growth with emergence of resistance: In phase 1, the cancer grows exponentially and resistance mutations lead to a treatment-resistant subpopulation. In phase 2, the sensitive population decreases while the resistant population continues increasing.

We investigate the emergence of resistance by dividing the population into treatment-sensitive and treatment-resistant cell types. We start with a single sensitive progenitor cell. When a sensitive cell divides, each daughter cells acquires resistance mutations with rate *ν*. Once a cell obtains a resistance mutations, it will be inherited by its offspring and the resistant clone is given new birth and death rates (fig. 1C). In total, we have 8 growth parameters: the birth and death rate of sensitive cells before treatment *b*_*s*,1_, *d*_*s*,1_ and during treatment *b*_*s*,2_, *d*_*s*,2_, and the same set of parameters again for resistant cells *b*_*r*,1_, *d*_*r*,1_ and *b*_*r*,2_, *d*_*r*,2_. We assume that resistant cells behave neutral in the absence of the drug (*b*_*s*,1_ = *b*_*r*,1_ and *d*_*s*,1_ = *d*_*r*,1_), sensitive cells decrease in number in the presence of treatment (*b*_*s*,2_ < *d*_*s*,2_) while resistant cells continue to increase (*b*_*r*,2_ > *d*_*r*,2_).

To model the accumulation of neutral mutations, we assume that in each division, each cell obtains *µ* mutations, where *µ* is a random number drawn from a Poisson distribution with mean *m*. Since each division comes with 2 daughter cells, there are on average 2*m* mutations generated per division. We assume infinite-sites such that each mutation is treated unique and mutation reversion is neglected.

We denote the stochastic total number of cells by *Z*_*0*_(*t*), its expectation by *N*(*t*) = *E*[*Z*_*0*_(*t*)] and add a tilde when conditioned on survival *Ñ*(*t*) = *E*[*Z*_*0*_(*t*)|*Z*_*0*_(*t*) > 0]. Next to the total population, subpopulations will follow stochastic growth too. We write *p*(*a* → *n, t*) for the probability that a (sub-)population has size *n* after time *t* when starting from size *a*. In its most general form, *p*(*a* → *n, t*) is a sum over min(*a, n*) terms that yield little analytic insight and is computationally expensive [12, 13]. Therefore, we synthesized exact and approximate results to balance computational efficiency and accuracy (Appendix A and *Methods*).

Some cancers, even if they have positive fitness, go extinct. If we observe a cancer it means that it survived genetic drift, which is mathematically incorporated by conditioning on survival. Given birth rate *b* and death rate *d*, we denote the probability for a population of size 1 to go extinct by time *t* with *α*(*t*). The extinction probability for a population of size *a* is then

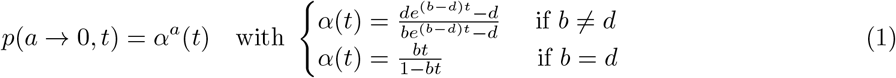

We can then write the average growth conditioned on survival by

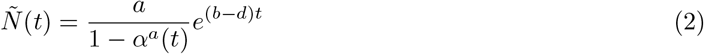

whereas the average growth including extinct trajectories is *N*(*t*) = *ae*^(*b*−*d*)*t*^ [14, 15].

Observations are made either at a fixed size or a fixed time. If we stop the stochastic process once the population reaches a fixed time, we have variable population size. The random size is fully characterized by *p*(*a* → *n, t*). Vice versa, if we stop at a fixed size, time will be a random variable. Conditioned on survival, the random time *T*_*N*_ to grow to size *N* for the first time when starting from a single cell is known to approximately follow a Gumbel distribution (ref. [15] and Appendix B) with expectation

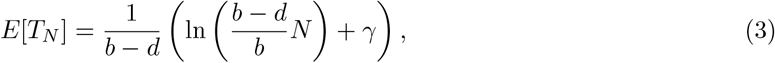

where *γ* ≈ 0.5772 is the Euler-Mascheroni constant. Both conditions have relevance in biological systems: In clinic, the starting time of the tumor is unknown and we observe the disease once it reaches detectable size. In contrast, in laboratory set-ups, experiments start and end at predefined times. It is possible to switch the conditioning of predictions between fixed-time and fixed-size (Appendix C). Using appropriate mapping between time and size, we find that the difference in the predictions between the two conditions is often small, in particular for large populations that behave nearly deterministically.

We investigate gITH of the cell population in terms of three summary statistics. First, we look at the site frequency spectrum (SFS). The SFS distributes mutations into classes 𝒮_*k*_ according to the number of cells *k* they occur in. The sizes of the classes 𝒮_*k*_ = |𝒮_*k*_| build a distribution 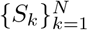 that we call the SFS. A site frequency corresponds to the abundance *k* of a mutation that resides at a particular site. Second, we study the total mutational burden (tMB) that we define as the number of unique mutations in the entire cancer and denote by *B*. The tMB counts mutations that are either clonal or subclonal and is related to the SFS by 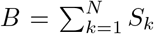. Third and last, we compute the single cell mutational burden (scMB) distribution. The scMB is the number of mutations in a single cell. The scMB distribution, 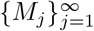, is defined by the number of cells, *M*_*j*_, with *j* mutations. In this sense, the scMB distribution distributes cells into classes according to the amount of mutations they carry.

Summary statistics can be computed from a single realization, and generally differ between different realizations. From this perspective, the SFS 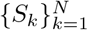 consists of *N* random variables, the total mutational burden *B* consists of 1 random variable, and the scMB distribution 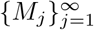 consists of infinitely many random variables. Mathematically speaking, the SFS and the scMB distribution are random distributions, i.e. distributions in which each value is a random variable. In the following, we are mainly interested in their expectations but also present some results of the variance between realisations. The latter corresponds to inter-tumour heterogeneity instead of intra-tumour heterogeneity.

## The site frequency spectrum

We link the SFS to the phylogenetic tree and derive an analytic formula for the expected SFS in homogeneous populations. We then apply this formula to obtain the expected SFS at the time of detection and after treatment accompanied with discussions on the properties and interpretations.

### Construction of the site frequency spectrum

Inspecting the phylogenetic tree in fig. 1A, we see that the number of mutations 𝒮_*k*_ that occur in *k* cells is obtained by summing over all mutations with branch width *k*. If we write the number of branches with width *k* as *W*_*k*_, label the branches with width *k* by *j* and the number of mutations on branch *j* by *µ*_*k,j*_, we have

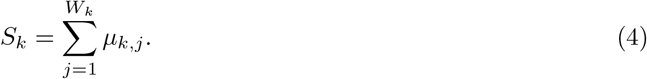

Since we assume infinite sites, the number of mutations *µ*_*k,j*_ are independent and by assumption follow a Poisson distribution, *µ*_*k,j*_ ∼ Poiss(*m*). We call the collection 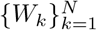 the branch width distribution (BWD), which is a summary statistic of the tree topology that emerged from the stochastic growth of the underlying cell population. Since eqn. (4) is a random sum, we can write the expectation and variance as

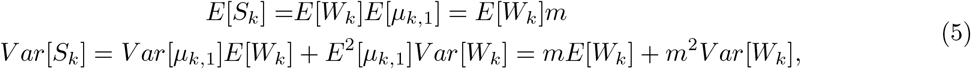

which disentangles the effects of stochastic growth and mutation accumulation. As we shall see in further proceedings, it is useful to separate between mutations that are preexisting to time *t*^′^ = 0 or newly emerging in a time interval [0, *t*]. We denote the number of newly emerging mutations found in *k* cells by 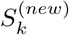, and the number of preexisting mutations by 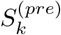. The total number of mutations 𝒮_*k*_ found in *k* cells is then 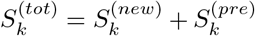. The following theorem characterizes the expected SFS of both contributions.

#### Theorem 1.

*Suppose the expected total population size over the time interval* [0, *t*] *is N*(*t*^′^) *with initial size N*_0_, *and the probability to grow from size x to y in time t is p*(*x* → *y, t*). *Then the expected SFS of newly emerging mutations conditioned on survival and fixed time t is*

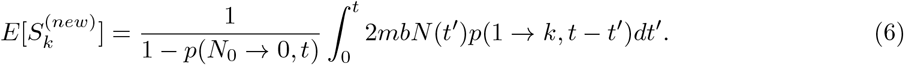

*Suppose the expected initial SFS at time t*^′^ = 0 *is* 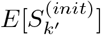, *then the expected SFS of pre-existing mutations conditioned on survival and fixed time t is*

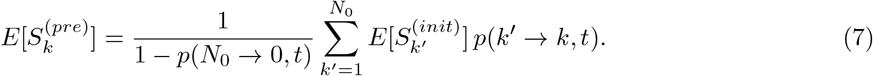

*Proof*. Appendix D.

For the proof of eqn. (6), we follow the derivation of ref. [16] who derived the result for *N*_0_ = 1. In both expressions, the first term is due to condition on survival of the entire population and can be dropped should one be interested in the expectation including extinction. Intuitively, eqn. (6) counts the number of mutations generated at time *t*^′^, conditions them to grow to size *k* and then sums over all mutations generated in the considered time interval. Eqn. (7) has similarities to a sampling formula. There are 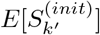 mutations of size *k*^′^ that are ”sampled” to size *k* with probability *p*(*k*^′^ → *k, t*). We do not strictly talk about sampling since we allow *k* > *k*^′^.

### The site frequency spectrum at detection

The expected SFS at detection, 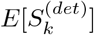, is computed by applying theorem 1 over the time interval [0, *t*_*d*_] starting from a single cell with no mutations (Appendix E). Following ref. [17] and [16], the resulting integral can be simplified to

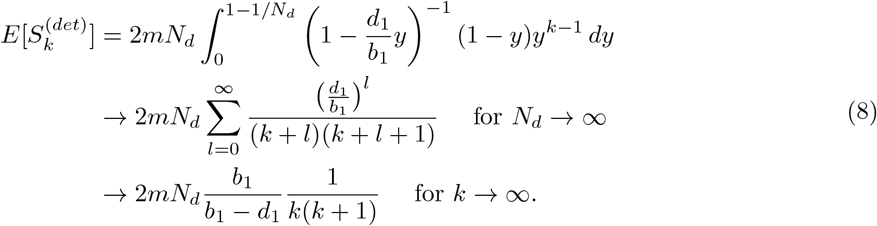

Here, *N*_*d*_ = *Ñ*(*t*_*d*_) is the expected population size at time *t*_*d*_ conditioned on survival. Although not the focus of this study, it is straightforward to include mutations accumulated before expansion that will be clonal in the entire cancer (Appendix E).

The prediction in eqn. (8) perfectly aligns with the average SFS obtained through computer simulations (fig. 2A). For large *k*, we have 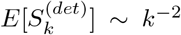, which is a well-known characteristic of the SFS in exponentially growing populations observable in bulk sequencing data of sufficient coverage [18, 5, 6]. Noticeably, single realisations of the stochastic process deviate significantly from the expected scaling for large site frequencies (2A, dots). These are mutations occurring in the first few generations of growth and have a high degree of stochasticity.

**Figure 2:**
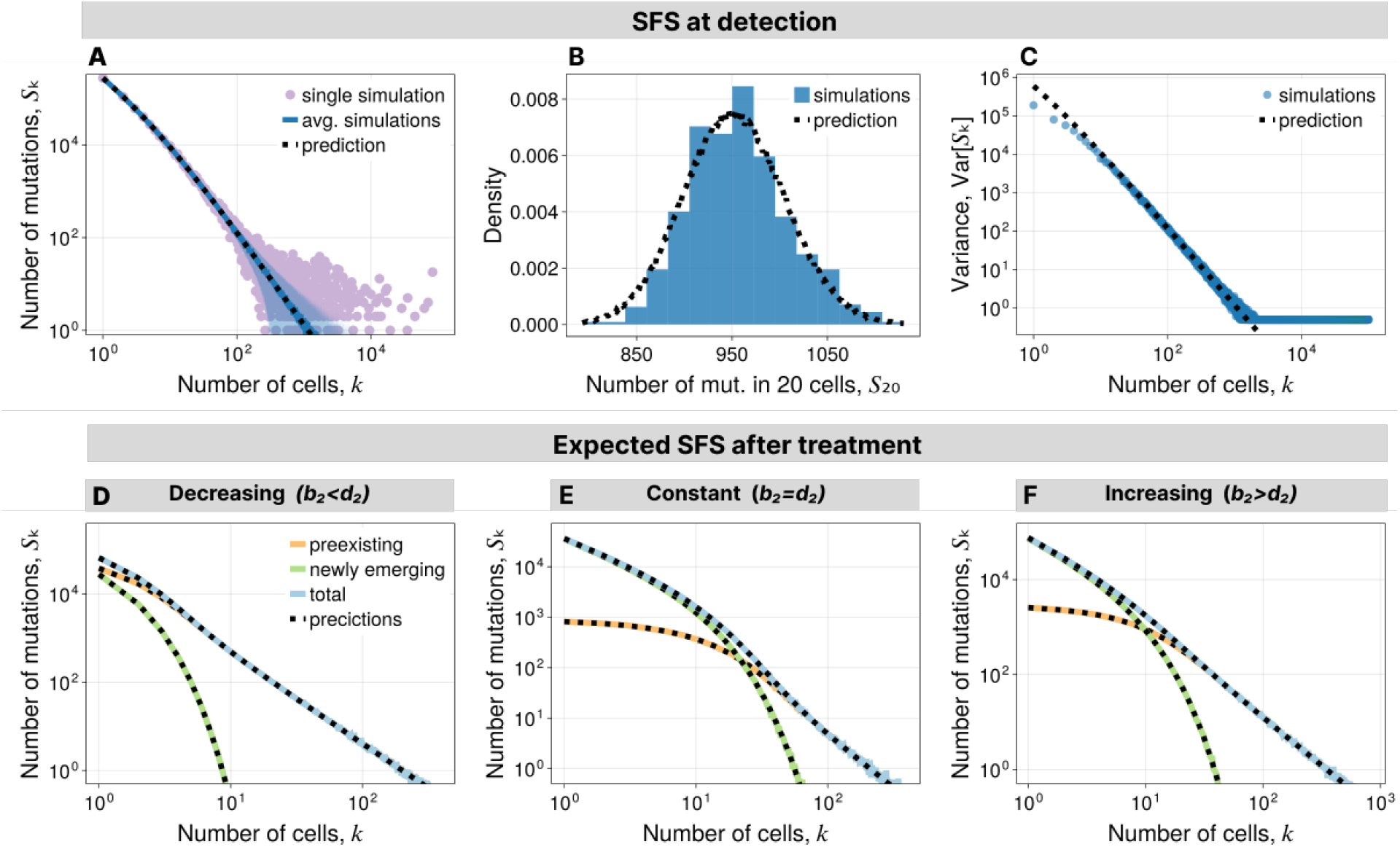
SFS in homogeneous populations. **A.-C.** SFS at the time of detection. **A**. Single simulation, average over many simulations with 1-sigma interval, prediction from eqn. (8). **B**. Distribution of S^20^ from simulations and prediction using a compound Poisson distribution. **C**. Variance in S^*k*^ from simulations and prediction from eqn. (10). **D.-F**. SFS after treatment. Average over many simulations and prediction from theorem 1 with explicit formulas in Appendix F. **D**. For decreasing population. **E**. For constant population. **F**. For increasing population. Parameters are in table S1.

Knowing the probability density 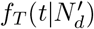 for the time *t* to reach fixed size 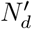 for the first time (Appendix B), we can change the conditioning from fixed time to fixed size (Appendix C). To avoid convergence issues, we consider the normalized SFS that we denote with a tilde 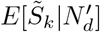. We change the conditioning by computing

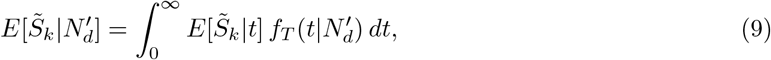

Using the large population size solution *N*_*d*_ → ∞ in eqn. (8), we show that the fixed-size expectation coincides with the fixed-time solution, 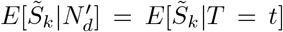 (Appendix E). This is in agreement with ref. [16] in which it is shown that the fixed-size expectation can be approximated by the fixed-time expectation for large *N*_*d*_.

The expected SFS *E*[𝒮_*k*_] after exponential growth is well studied and routinely observed in cancer genetic data. However, the variance *Var*[𝒮_*k*_] of the SFS under exponential growth remains to the best of our knowledge unknown. In the pure-birth process, and within the fixed-size limit, we observe in simulations that *W*_*k*_ are approximately Poisson distributed such that 𝒮_*k*_ is described by a compound Poisson process (fig. 2B). This suggests a heuristic expression for the variance

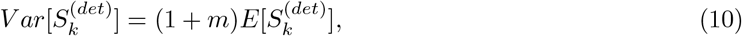

that shows good agreement with simulations of the pure-birth process on fixed size (fig. 2C). There are deviations for very small site frequencies *k* = 1, 2, 3 To better understand this, we point out that when conditioning on growth to size *N*_*d*_, there are exactly *N*_*d*_ branches, which are simultaneously leaves such that 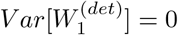 and thus 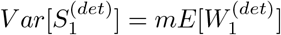.

Interestingly, eqn. (10) suggests that the variance in the SFS carries information on the mutation rate *m*. This can be leveraged to infer the mutation rate *m* from sequencing data, possibly at different stages of tumour growth.

### The site frequency spectrum after treatment

We compute the SFS after homogeneous treatment response, and distinguish between decreasing, constant and continued increasing cell populations (fig. 1B). To apply theorem 1, we assume that the cancer has initially fixed size 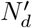 and expected SFS at detection, 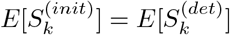. For analytical purposes, we approximate the fixed-size expected SFS with the fixed-time expected SFS in eqn. (8). Together with the expected growth 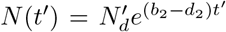, the treatment time interval [0, *t*_*f*_] and probabilities *p*(*a* → *n, t*) (Appendix A), we can readily compute the expected SFS of newly emerging and preexisting mutations. We validate the general analytic predictions computed in Appendix F with simulations showing perfect alignment (fig. 2D-F). We proceed with highlighting some approximations and observations.

First, we observe that newly emerging mutations are restricted to small site frequencies, and thus the tail is determined by preexisting mutations. In shrinking populations, newly emerging mutations are prone to extinction and only survive short amounts of time and thus have almost negligible impact on the SFS. In contrast, for constant or increasing tumour populations, newly emerging mutations dominate the SFS at low frequencies whereas preexisting mutations continue to dominate the SFS at high frequencies (fig. 2D-F).

For decreasing populations, we obtain a simple formula in the case of the pure-death process. To show this, we adapt the sampling process from theorem 3 in ref. [15] (Appendix G). Taking *b*_2_ = 0 and sufficiently large *k*, we find

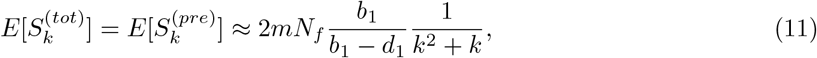

where *N*_*f*_ = *Ñ*(*t*_*f*_) is the expected population size at time *t*_*f*_ conditioned on survival. There are no newly emerging mutations during treatment since *b*_2_ = 0, and the *k*^−2^ tail of neutral mutations from the SFS at detection is maintained.

As *b*_2_ → *d*_2_, the importance of genetic drift increases, and if *b*_2_ = *d*_2_, the SFS of newly emerging mutations takes the form

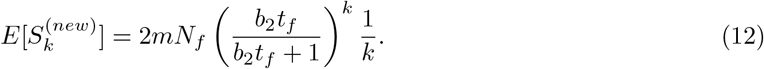

As *t*_*f*_ → ∞, all preexisting mutations will be either extinct or fixed in the population such that only newly emerging mutations contribute to gITH. From eqn. (12), we can see that the SFS scales with *k*^−1^ in agreement with previously reported results for the SFS in constant populations at equilibrium [19, 20, 16].

### Testing homogeneity

Theorem 1 provides us predictions for the SFS at detection and after treatment assuming that all cancer cells have the same growth parameters. In the spirit of neutral theory [21, 22], we can take this prediction as null hypothesis of homogeneity that can be compared to sequencing data [18]. If the population is homogeneous, we expect a good fit between the data and the prediction. If the population is heterogeneous, we expect to see deviations between data and prediction.

Sequencing data are noisy and mutations that occur in less than 1% of the cell population cannot reliably be observed. This restricts our test on the observable mutations occurring in at least 1% of the population 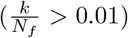. However, considering large site frequencies dramatically simplifies the homogeneity test in the treatment scenario. From simulated data, we observe that the SFS in mutations above 1% of the population follow the power law S_*k*_ ∼ *k*^−2^ known from exponentially growing tumors (fig. S2). We support this observations by arguing that the observable SFS for biologically feasible parameters consists of large site frequencies that behave nearly deterministic (Appendix G). Remarkably, this holds true for all three considered cases of homogeneous growth, although deviations may occur for small population that are kept constant for long times [9]. We conclude that testing homogeneity can be achieved by evaluating whether the SFS follows the expected scaling 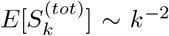 as demonstrated in prior analyses of the SFS at detection [18, 5].

## The total mutational burden

Analogous to the SFS, we link the tMB to the topology of the phylogenetic tree, and provide a general formula to compute the expected tMB, which we then apply to homogeneous cancer cell populations before and after treatment. We connect the tMB to the risk of resistance and discuss its relevance for therapeutic interventions.

### Construction of the total mutational burden

Looking at the phylogenetic tree, we denote the number of nodes including leaves with at least one living descendent by *R*, and label them with *i* = 1, 2, 3, …, *R*. Each node comes with *µ*_*i*_ unique mutations such that the tMB is

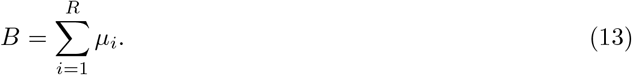

Here, *R* and *µ*_*i*_ are random numbers and *µ*_*i*_ ∼ Poiss(*m*) are independent such that

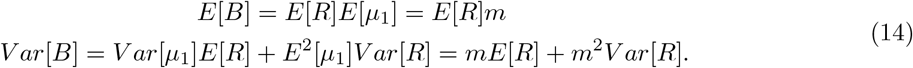

We separate again between newly emerging and preexisting mutations such that the total tMB is the sum of the two contributions, *B*^(*tot*)^ = *B*^(*new*)^ +*B*^(*pre*)^ that are characterized with the following theorem.

#### Theorem 2.

*Suppose the expected total population size over the time interval* [0, *t*] *is N*(*t*^′^) *with initial size N*_0_, *and the probability to grow from size x to y in time t is p*(*x* → *y, t*). *Then the expected tMB of newly emerging mutations conditioned on survival and time t is*

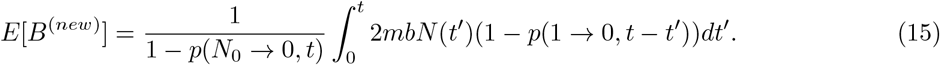

*Suppose the expected initial SFS at time t*^′^ = 0 *is* 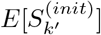, *then the expected tMB of preexisting mutations conditioned on survival and time t is*

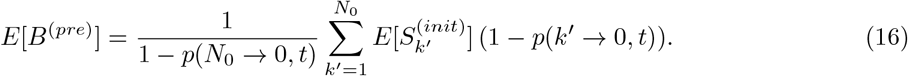

*Proof*. Appendix H.

The expected tMB can be obtained from theorem 1 by noting that 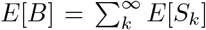. However, the expressions for the expected SFS after treatment are complicated and theorem 2 then provides more straightforward calculations.

### The total mutational burden at detection

Following theorem 2, the expected tMB at detection is

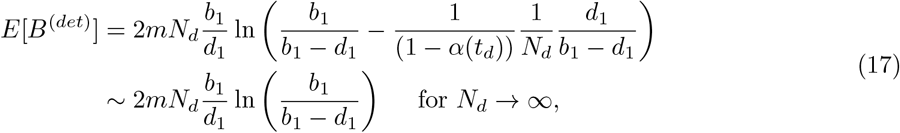

where *N*_*d*_ = *Ñ*(*t*_*d*_) is the expected size at fixed time *t*_*d*_. The expression coincides with the expression for the tMB in ref. [16], in which the expected tMB was computed by summing over the SFS. We focus on mutations that occur during growth and treatment. Should one be interested in mutations preexisting in the ancestor cell, those will be shared by the entire population and can be added as a constant. Detailed calculations are in Appendix I.

From eqn. (17), it is apparent that increasing the death rate increases the tMB if measured at the same population size. This is expected, as an increased death rate implies more cell divisions to reach the same population size. However, a considerable fraction of mutations are at low frequencies and will go extinct even if the cell population continues increasing.

In the pure-birth process, the tMB is simply *E*[*B*^(*det*)^] ≈ 2*mN*_*d*_. To grow from 1 cell to *N*_*d*_ cells, the population undergoes *N*_*d*_ − 1 divisions, where each daughter cells obtains a random number of mutations. Thus, the tMB is the sum of 2 × (*N*_*d*_ − 1) independent random variables that are Poisson distributed with mean *m*. Approximating *N*_*d*_ − 1 ≈ *N*_*d*_, we have *B*^(*det*)^ ∼ Poiss(2*mN*_*d*_), which agrees with stochastic simulations conditioned on fixed size (fig. 3A). For observations made on a fixed time, the varying number of cells yields an additional layer of stochasticity. Also, if *d*_1_ > 0, there are different growth trajectories to reach size *N*_*d*_. From eqn. (14), it is clear that in either of the two cases *Var*[*B*^(*det*)^] > *E*[*B*^(*det*)^] because the variance in division counts is positive, *Var*[*R*^(*det*)^] > 0, implying an increased genetic heterogeneity between tumours (fig. 3B-C).

**Figure 3:**
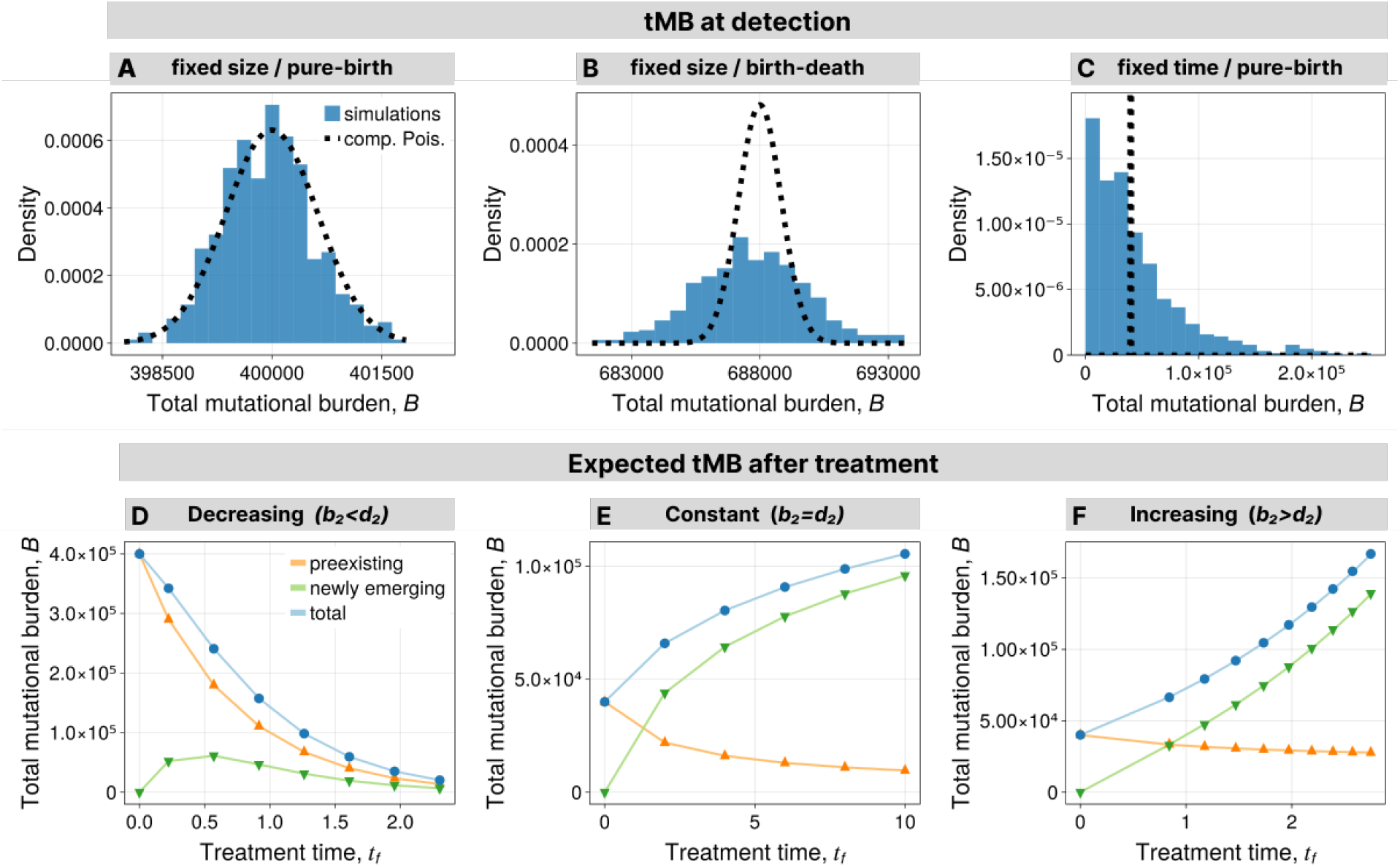
tMB in homogeneous populations. **A.-C.** Distribution of tMB at detection from simulations compared to compound Poisson distribution with mean given by eqn. (17). **A**. Pure-birth process conditioned on fixed size. **B**. Birth-death process conditioned on fixed size. **C**. Pure-birth process conditioned on fixed time. **D.-F**. Average tMB after treatment from simulations (points and triangles) and predictions (solid lines) using theorem 2 with explicit formulas in Appendix J. **D**. For decreasing population. **E**. For constant population. **F**. For increasing population. Parameters are in table S2.

### The total mutational burden after treatment

We compute the expected tMB after homogeneous treatment response using theorem 2. We assume initial size 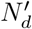, expected SFS 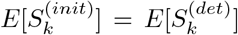 approximated by eqn. (8), and treatment interval [0, *t*_*f*_]. The tMB of mutations emerging during treatment is solved exactly. For decreasing or increasing population (i.e. *b*_2_≠ *d*_2_), it is

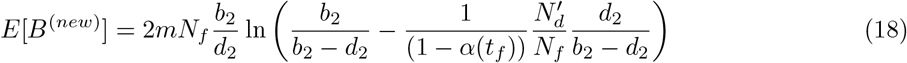

that simplifies to eqn. (17) for 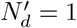. For the constant population (*b*_2_ = *d*_2_), it is

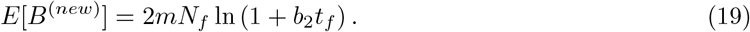

The expression for mutations preexisting to treatment is more involved due to summation over the initial SFS. However, we find a simple approximation for *d*_1_ = 0, which we heuristically extend for general *d*_1_ > 0 that reads

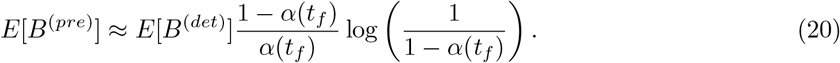

The separation between *b*_2_ = *d*_2_ and *b*_2_≠ *d*_2_ is made through *α*(*t*_*f*_) defined in eqn. (1). The approximation performs very well in all three cases with small deviations for increasing *d*_1_, in which case the approximation yields an useful upper bound (fig. S3).

In fig. 3D-F, we show the alignment of the exact theoretical predictions with stochastic simulations. It is worth noting that most mutations contributing to tMB are at low frequencies. After exponential growth, it is estimated that 50% of unique mutations occur in only 1 cell [16]. After treatment, in particular if the population remains approximately constant, this asymmetry is less extreme [9]. Still, the large majority of mutations remain at low frequencies.

### Relationship to genetic resistance against therapy

The tMB is linked to the risk of resistance. An increasing number of mutations comes with an increased chance of genetic resistance mechanisms, which is confirmed in clinical data for various cancer types and treatments [4, 11]. Mathematically, the tMB is connected to the probability of having a resistant subpopulation through a probability *p*_*r*_ that a specific mutation leads to resistance. Given the tMB is *B*(*t*) at time *t* the probability that none of those has led to resistance is Pr(sensitive) = (1 − *p*_r_)^*B*(*t*)^. Although this picture is oversimplified, it makes clear that reducing the tMB generally decreases the risk of resistance. Notable exceptions are immunotherapies, which rely on the accumulation of neoantigens to be effective. In this case, an increased tMB is a double-edged sword: Higher tMB comes with an increased treatment efficiency due to a higher load of neoantigens necessary for the immune system to detect the cancer cells as invaders but also increases the chance of resistance mechanisms such as immune escape [23, 24, 25].

In fig. 3D-F, we show that mutations arising during treatment contribute significantly and often dominate the tMB shortly after treatment initiation. Especially in the constant population scenario, over time, preexisting mutations are almost fully depleted and most mutations will have arisen during treatment. This is an important observation, especially for the planning of possible adaptive clinical trials that are already ongoing or in preparation [26, 27]. A constant population during treatment approximates adaptive therapy cycles in the limit of small cycle sizes. Our calculations suggest that drugs suppressing the accumulation of new mutations - if feasible - would significantly reduce the risk of novel acquired genetic resistance in such patients, while at the same time weeding out preexisting genetic diversity.

There also has been a long debate if treatment resistance is mostly caused by selection on preexisting genetic diversity or resistance is acquired during periods of treatment [28]. These questions are not only related to cancer therapies but similarly can be asked for antibiotic or antiviral therapies [29, 30]. As we show in fig. 3D-F, there is not a single correct answer, but the dominant mechanism does both depend on the timing and duration of therapy and the population dynamics of the treated population. In case of a shrinking population, diversity is for the most part dominated by preexisting mutations. In contrast, in constant or growing populations, genetic diversity is very fast dominated by newly acquired mutations. These observations should be taken into consideration in future planning of multi-drug combination or adaptive cancer therapies and possible underlying mechanisms of failed clinical trials.

## The single cell mutational burden distribution

The single cell mutational burden (scMB) distribution has started recently to attract attention. Whole genome single-cell mutational burden information across different normal human tissues and ages has become available recently [1] and theoretical models of single cell mutational burdens during homeostasis have shown to complement information encoded in the SFS [9]. We extend on these considerations and discuss the scMB distribution and its properties under scenarios of cancer growth and during treatment focusing on homogeneous populations.

### Construction of the single cell mutational burden distribution

The scMB distribution *M*_*j*_ is linked to the phylogenetic tree through the divisional distributions, which counts the number of cells *D*_*l*_ with *l* divisions. In terms of the tree topology, *l* is the leaf depth that is the distance, i.e. number of nodes, from the leaf to the root (fig. 1A). The distribution 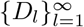 partitions the cell population into classes of cells according the number of divisions. Picking a cell with label *m* that has undergone *l* divisions, labelling its divisions by *n* = 1, 2, …, *l* and noting that on each division the cell accumulated *µ*_*m,n*_ mutations, we can write the scMB distribution in term of the divisional distribution. Namely, we have

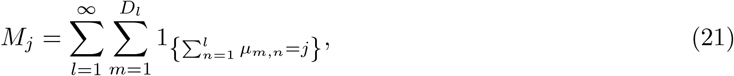

where 1_{*x*=*y*}_ is an indicator function that is 1 if *x* = *y* and 0 otherwise. The following theorem characterizes the expected scMB distribution.

#### Theorem 3.

*Given population size N, the expected scMB distribution is expressed in terms of the epected divisional distribution E*[*D*_*l*_] *by*

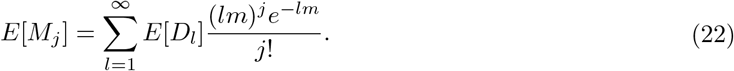

*Starting with N*_0_ *cells that have undergone zero divisions, the mean-field solution for the expected divisional distribution for a population of size N at time t is*

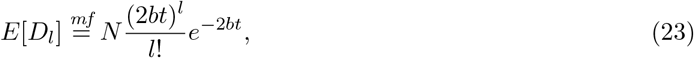

*such that the probability mass* 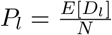 *follows a Poisson distribution with mean* 2*b*t.

*Proof*. Appendix K.

For exponentially growing populations, eqn. (22) was also recently reported in ref. [31]. Eqn. (23) was derived in ref. [32] and [5]. Furthermore, it has been shown that the divisional distribution is normal distributed as time goes to infinity [33, 34, 35] and some exact expressions with higher complexity were obtained [36, 37]. Since the mean-field solution shows good agreement with simulations up to a small correction for small population size, we proceed with the much simpler mean-field solution.

The average number of divisions for a single cell underlies an inspection bias [36]. Consider a population that has lived for time *t* with birth rate *b*. Picking a leaf of the phylogenetic tree at random and tracing it back to the root, one may naively estimate that the leaf has undergone *bt* divisions on average. Yet, eqn. (23) predicts an average division number of 2*bt*. To understand the increased number of divisions, note that some cells proliferate more than other by chance. Cells with more divisions have a larger number of offspring and contribute more to the average number of divisions in the observed population.

### The single cell mutational burden distribution at detection

Following theorem 3, we first obtain the divisional distribution from eqn. (23) and then the scMB distribution from eqn. (22) for a cell populations starting with 1 cell growing to *N*_*d*_ cells. Comparison between the analytic prediction and simulations revealed a small constant offset of the mean. To resolve this discrepancy, we looked at the average number of divisions per cell defined by 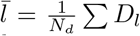. Rigor treatment of the pure-birth process shows that the expected average number of divisions is given by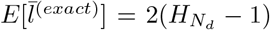. From this it follows that the mean-field solution obtained through eqn. (23) differs from the exact solution by a constant 2 when conditioned on fixed size or ln(4) when conditioned on fixed time (Appendix L). Focusing on fixed-size observations, we integrate the correction constant into the divisional distribution setting

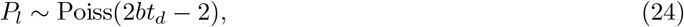

where *t*_*d*_ is defined as the mean time to reach size *N*_*d*_ for the first time (see Appendix B). The adjusted scMB distribution shows excellent agreement with simulations of the pure-birth process (fig. 4A), and good but not perfect agreement when deaths are included (fig. S5). The scMB distribution of single realisation has the same shape as the expected distribution, although it comes with a small random shift of the mean. These shifts are caused by stochastic effects at small population sizes leading to mutations shared by a large fraction of the population.

**Figure 4:**
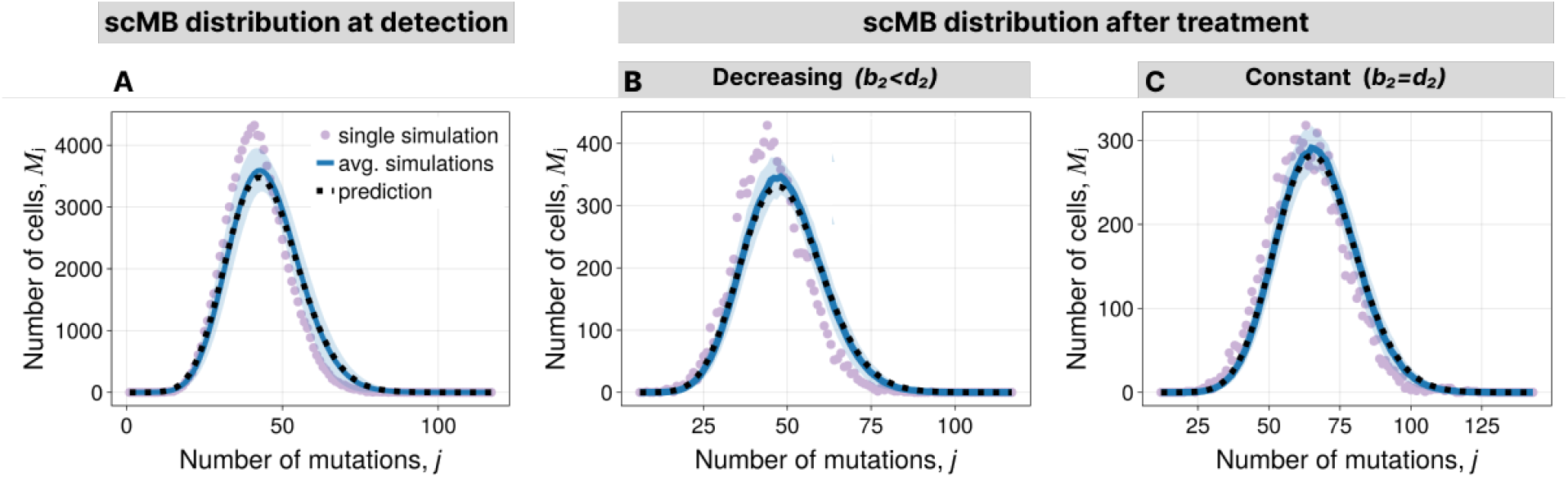
**A.** scMB distribution at detection for the pure-birth process. Average over many simulations with the 1-sigma interval, single realization, and the prediction from theorem 3 with divisional distribution as in eqn. (24). **B.-C**. scMB distribution after treatment. Average over many simulations with the 1-sigma interval, single simulation, and the prediction from theorem 3 with divisional distribution as in eqn. (25). **B**. For decreasing population. **C**. For constant population. Parameters are in table S3.

### The single cell mutational burden distribution after treatment

For homogeneous treatment response, the analytic prediction for decreasing, constant and increasing populations under treatment is the same. As long as the final population size is large, i.e. *N*_*f*_ ≫ 1, small population effects can be neglected such that the average number of divisions obtained during treatment time 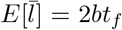. Those divisions are again Poisson distributed and together with divisions before treatment, we have

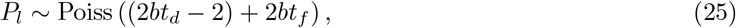

where 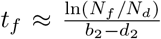. We put *E*[*D*_*l*_] = *N*_*f*_ *P*_*l*_ into eqn. (22) to obtain the scMB distribution showing good alignment with simulations (fig. 4B-C, fig. S6).

The average scMB increases linear in time (fig S4), in concordance with experimental observations of healthy human tissues [2]. Whereas this is also true for some limiting cases for the tMB normalized over the population size, the tMB shows in general more complex non-linear behaviour (fig. 3D-F). Similarly to the observable SFS, the scMB distribution does not change its shape after homogeneous treatment response yielding a null model to test homogeneity. In contrast to the SFS, the scMB distribution differs from the null hypothesis only for heterogeneity in birth rates, i.e. resistant cells have different birth rates than sensitive cells. Furthermore, the scMB distributions allows decoupling mutation rate and birth rate by studying its variance [9].

## Genetic intra-tumour heterogeneity upon emerging resistance

So far, we discussed properties of gITH in cell populations with homogeneous treatment response. For certain treatment regimes such as chemo or radio therapy in early stage disease, this seems a reasonable null scenario. However, in other cases, emergence of resistant subpopulations is a major problem. Genetic resistance is well documented in many targeted cancer therapies, where specific point mutations induce resistance [38, 39]. Resistance mutations give rise to resistant clones leading to a cell population, in which some cells are less or not affected by treatment (fig. 1C). Building up on the results for homogeneous populations, we first study the size distribution for resistant clones, followed by investigations of the SFS and scMB distribution.

### The resistant clone size distribution

We distinguish between mutations without effect accumulated at rate *m* and mutations that cause resistance accumulated at rate *ν* per cell and division. We denote the number of resistant clones of size *κ*, by Θ_*κ*_ and call 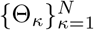 the clone size distribution (CSD). The CSD and the SFS have a conceptional difference: the CSD is not nested, i.e. clones are not overlapping, whereas the SFS allows nested lineages. However, since *ν* is very small, resistance mutations have negligible effect on the growth dynamics of sensitive cells from which the resistant clones arise. As a consequence, we can approximate the CSD by adapting theorem 1 (Appendix M).

At detection, the expected CSD is

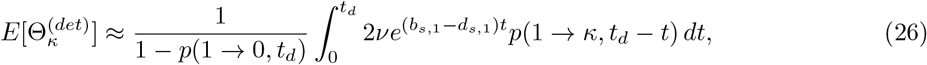

where the term for conditioning on survival 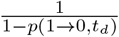 is appropriately parameterized with growth parameters for sensitive cells *b*_*s*,1_, *d*_*s*,1_ while the term *p*(1 → *κ, t*_*d*_ − *t*) must be parameterized with growth parameters for resistant cells *b*_*r*,1_, *d*_*r*,1_. Assuming that the growth parameters of sensitive and resistant cells are the same in the absence of treatment, the CSD takes the same form as the SFS in homogeneous populations in eqn. (8). Focusing on the limit *N*_*d*_ → ∞ and writing *b*_1_ := *b*_*s*,1_ = *b*_*r*,1_ and *d*_1_ := *d*_*s*,1_ = *d*_*r*,1_, we have

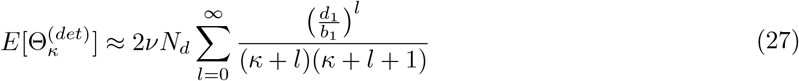

This is not surprising, given above assumptions, resistance inferring mutations are neutral before treatment.

Next, we consider the CSD after treatment time *t*_*f*_. As initial condition, we assume fixed size 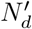, and initial 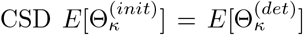. We adapt theorem 1 and obtain the expected CSD of clones that emerged before and after detection giving us

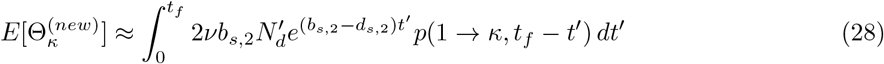

and

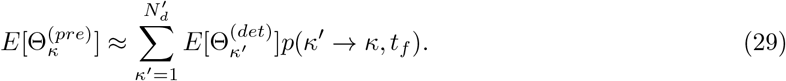

In eqn. (28), the relevant mutation generation is caused by sensitive cells 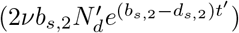. In both eqn.(28) and (29), the growth of clone sizes *p*(*a* → *n, t*) is parameterized for resistance cells *b*_*r*,2_ and *d*_*r*,2_.

### Subdominance of the largest resistant clone

In an exponentially growing population, the first mutant clone has a large advantage as all later arriving clones need to catch up to become dominant. We thus asked whether the first resistant clone always dominates all other later arising resistant clones. Here, we show that this is not the case. We focus on detection since a dominating resistant clone at detection also dominates after treatment.

First, we find that the first arriving clone is not always the largest. We derived the probability densities for the arrival time *T*_1_ of the first clone and the second clone *T*_2_ given that *T*_1_ = *t*_1_ and survival of drift (Appendix N). The long-term growth of the clones can be written as 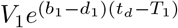 and 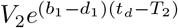 where *V*_1_, *V*_2_ are exponentially distributed with mean 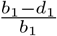 (Theorem 1 in ref. [15]). The difference in their size at time *t*_*d*_ is then

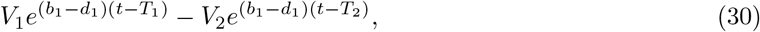

which we sample and illustrate in fig. 5A. One can see that a considerable fraction is in the negative range (illustrated in red). This means that the second clone can outgrow the first clone purely by drift. These effects would become more pronounced for possible fitness distributions of resistant clones.

**Figure 5:**
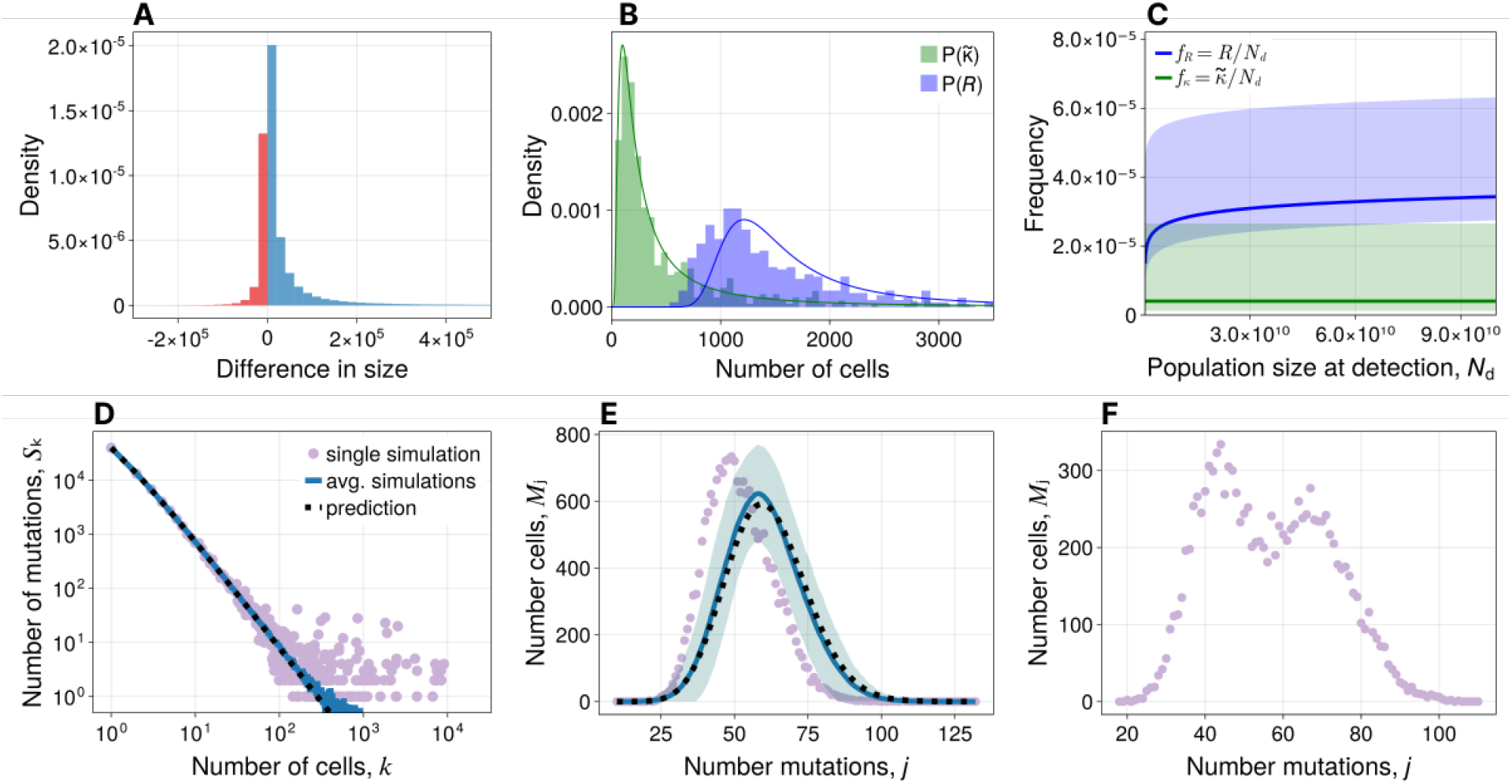
**A.** Difference in size between the first and second resistant clone at detection given by eqn. (30). Negative differences are shown in red, and positive differences in blue. **B**. Size distributions for the largest resistant clone and the resistant subpopulation at the time of detection. Simulations as histograms, and predictions as solid lines given by eqn. (31) and (32). **C**. Distribution of frequencies for the largest resistant clone and the resistant subpopulation at the time of detection. The median frequency is shown as a solid line with shadow ranging from the 10% to the 90% quantiles. **D**. SFS after treatment upon emergence of resistance from simulations and prediction of the compound tail in eqn. (34). Parameters are set such that all sensitive cells are extinct. **E**. scMB distribution upon emergence of resistance in which birth rates remain unaffected. For the prediction, we used treatment time *t*^*f*^ measured from one simulation. **F**. scMB distribution for a single population upon emergence of resistance with change of birth rates. Parameters are in table S4.

Next, we considered the largest clone at time *t*_*d*_ instead of the first arriving clone. Interpreting the CSD as a probability distribution for clone sizes, we can use methods from order statistics [40] and find an approximate probability density for the size of the largest resistant clone 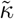 that reads

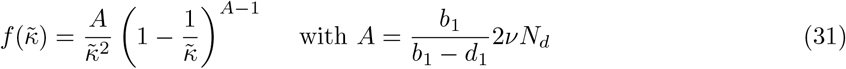

and is valid for *νN*_*d*_ ≫ 1. The derivation is presented in Appendix N and we validated our approximation with simulations (fig. 5B).

We compare the size of the largest resistant clone to the total number of resistant cells. Given small mutation rate *ν* but large population sizes *N*_*d*_ and *νN*_*d*_ ≫ 1, the probability of having *R* resistant cells at the time of detection is described by a Landau distribution [41, 42]. We have

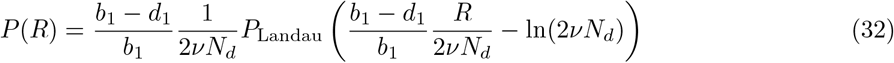

that also shows good agreement with simulations but with a small offset (fig. 5B).

Both distributions have a fat tail that decreases with exponent −2. The mean and the median of the largest clones 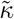 are always below the mean and median of *R*. Interestingly, the median fraction of all resistant cells is weakly increasing with detection size *N*_*d*_ and is predicted to be around 3.0 × 10^−5^ for biologically feasible parameter values. In contrast, the median size of the largest resistant clone is approximately constant for different detection sizes and is around 4.0 × 10^−6^ (fig. 5C). This is reasonable as newly resistant cells continue to emerge from sensitive cells during growth and thus increase the fraction of resistant cells as the tumour grows. We conclude that the largest clone may make up a significant fraction but probably not a dominating fraction of the resistant subpopulation.

### The site frequency spectrum

We assume resistant cells to be neutral in the absence of treatment such that the expected SFS at detection is unchanged given by eqn. (8). In the following, we focus on the SFS after treatment. We separate between neutral mutations that emerged in sensitive cells 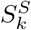 and mutations that emerged in resistant cells 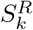. We label *K* resistant clones with *i* = 1, 2, …, *K* such that

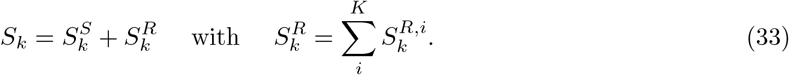

By definition, 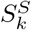 combines mutations in sensitive cells and mutations in resistant cells acquired in sensitive ancestors.

If resistance mutations lead to full resistance, i.e. *b*_*r*_ := *b*_*r*,1_ = *b*_*r*,2_ and *d*_*r*_ := *d*_*r*,1_ = *d*_*r*,2_, then the compound SFS of all resistant clones takes again the known form

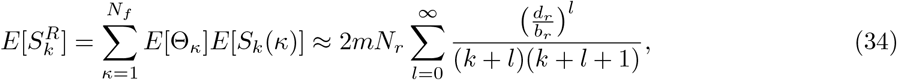

where *N*_*r*_ is the number of resistant cells, *E*[S_*k*_(*κ*)] is the SFS of an exponentially growing cell population at size *κ*. The calculations are presented in Appendix O and alignment with simulations is shown in fig. 5D.

To make progress on 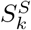, we assume sufficiently long treatment times such that all sensitive cells go extinct. Then, 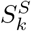 consists of mutations in resistant cells that are clonal in at least one resistant clone. If we denote 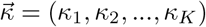 as a vector of clone sizes *κ*_*i*_ with indices *i* ∈ *X*, we have

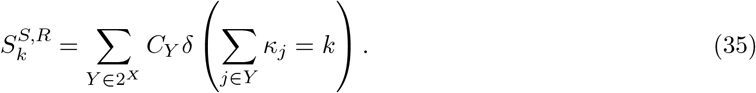

Here, the first sum goes over all possible combinations *Y* in the set of indices *X* that is the power set 2^*X*^, and C_*Y*_ is the number of mutations shared between the chosen clones. Those mutations will arise as peaks in the SFS (fig. 5D and S7). The heights and the locations of the peaks carry information on the timing and selection of resistant clones. However, a more in-depth understanding of the peaks requires analysing the relatedness between clones and goes beyond the scope of this study.

### The single cell mutational burden

From our previous investigations, we know that the scMB is not affected by the death rate but only the birth rate. Thus, if the birth rate is unaffected in the selection process then the scMB distribution remains the shape described in theorem 3, which we confirmed with simulations (fig. 5E). However, if the birth rate of the resistant cell lines differs from the sensitive cells then the scMB distribution will differ too (fig. 5F).

We write the scMB distribution of the resistant population as 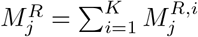, where 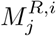 is the scMB distribution of an individual clone with label *i*. Together with the sensitive cells, the total expected scMB distribution is

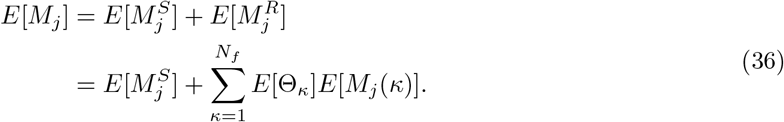

The total scMB distribution consists of overlapping scMB distribution of sensitive cells and resistant clones. If the birth rate changes in resistant cells during treatment but is the same among all resistant cells, the majority of resistant cells consist of clones that arose before treatment with the same expected scMB leading to a combined single peak of resistant cells next to the peak of sensitive cells (fig. 5F). If the birth rate of resistant clones changes already before treatment, the clones will have different expected scMB leading to many overlying distributions.

## Discussion

We used stochastic processes to investigate expected patterns of genetic heterogeneity in cancer before and after treatment. We discussed three summary statistics of gITH (SFS, tMB, scMB) for different treatment scenarios. For homogeneous treatment response, we found that the SFS keeps its *k*^−2^ power law tail regardless whether the cancer is shrinking, approximately constant or growing, and the scMB distribution is described by two entangled Poisson processes. Those predictions build the basis for possible homogeneity tests, and can provide insights into the mechanisms underlying resistance evolution. Similar usage of a null hypothesis of the SFS has led to biological insights into selection processes of healthy tissues [43] and cancer before treatment [18]. We further characterised the non-linear behaviour of the tMB. Notably, the tMB relates to the probability of having resistant cells and the scMB may be interpreted as scMB of antigens important for the response to immunotherapies.

Next, we investigated the effect of treatment resistance on gITH assuming a simple model of genetic resistance reasonable for treatment of late-stage cancers with targeted therapies that often show resistance due to specific point mutations [38, 39]. We found that the resistant subpopulation will consist of many resistant clones that arose from sensitive cells. The largest resistant clone likely makes up a considerable but not dominating fraction of the total resistant subpopulation. We showed that resistant clones leave traces in the SFS, and change the scMB distribution only if birth rates are affected.

For generality and tractability, here we used a minimal set of assumptions when defining our theoretical framework. Data generation often comes with additional complexities that must be incorporated before our results can be readily compared to the data. First and most important, sequencing comes with sampling effects as well as sequencing noise. However, these can be taken into account in both stochastic simulations and data analysis [9, 5]. Furthermore, there are possible biological complexities including (i) the impact of spatial structure [44] (ii) complex selection processes, e.g. varying levels of treatment response among resistant clones [45] (iii) the impact of the tumour-microenvironment and other non-genetic determinants [46]. These effects may contribute to evolutionary paths of individual tumours. As such, our results provide testable null hypotheses of which additional complexities can be tested. We hope that our results will set the stage for further theoretical and data-driven investigations.

## Methods

Computer simulations were implemented in the Julia programming language and are based on the Gillespie algorithm [47, 48]. We adapted the simulation framework developed for ref. [49]. The cell population is saved in a tree structure. The root of the tree represents the ancestor cell and the leaves represent living cells. In case of a death event the chosen leaf is removed from the tree. In case of a birth event two new leaves are created and each carries the number of mutations drawn from a Poisson distribution. Then, each node that is not a leave represents an ancestor of a currently living cell.

To obtain the SFS, we iterate over all nodes and count the number of mutations unique to this node and its ancestors. If the node with label *i* has *µ*_*i*_ mutations and leads to *k* leaves, we increase S_*k*_ by *µ*_*i*_. After iterating over all nodes, we have the SFS.

For the scMB distribution, we iterate over all leaves. On top of the mutation of the leaf itself, we add its parent’s mutations, then its parent’s parent’s mutations and go on until we reach the ancestor cell. The final sum is the scMB of this leaf. After iteration over all leaves, we have the scMB distribution.

Numerical solutions of integrals were computed using the HCubature module in the Julia programming language [50] that is based on an adaptive algorithm [51]. Integrals were computed up to a relative tolerance of 10^−3^.

Mathematical analysis was supported using *Wolfram Mathematica* [52].

We balance efficiency and accuracy for the computation of the probability mass function of the birth-death process *p*(*a* → *n, t*). We always use exact solutions for the pure-birth process, pure-death process or when the population start from a single cell [53]. Otherwise, we use exact solutions written as computationally stable sum [12] for min(*a, n*) ≥ 50 and use the (unnormalized) saddlepoint-approximation method solution otherwise [13]. Details of the approximations are described in Appendix A.

Random samples for random variables *T*_1_ and *T*_2_ were generated using inverse transformation sampling [54]. Therefore, a random number *u* is drawn from a uniform distribution in (0, 1) and is then transformed into a random number *x* with cumulative density function *F*_*X*_ (*x*) by plugging the uniform random number into the inverse distribution function 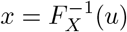.

We fit the exponent *γ* of the power-law S_*k*_ ∼ *k*^*γ*^ to analyse the SFS obtained from simulated data. Therefore, we use a maximum likelihood estimator for discrete power law distributions from ref. [55]. Adapted for our purposes, we have

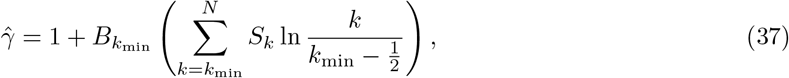

where *k*_min_ defines the smallest site frequency used for the fitting, 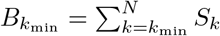 is the number of mutations that occur in at least *k*_min_ mutations and *N* is the population size.

## Acknowledgement

AS was supported by the European Union’s Horizon 2020 research and innovation program under the Marie Sklodowska-Curie EvoGamesPlus grant agreement No. 955708. BW was supported by a Barts Charity Lectureship (grant no. MGU045) and a UKRI Future Leaders Fellowship (grant no. MR/V02342X/1). We thank Nathaniel Mon Père and Christo Morison for helpful discussions at different stages of the manuscript.

## Code availability

For the simulations, we adapted a package that was written for ref. [49] and is available at https://github.com/jessierenton/SomaticEvolution.jl.

The adapted package is available at https://github.com/alexsteininfo/TreeStatistics.jl and code to run and analyse the simulations is available at https://github.com/alexsteininfo/GITH-Treatment-Patterns.

## Appendix

### A Probability mass functions of the birth-death process

Given constant rates for birth *b* and death *d*, and starting with *a* cells, what is the probability *p*(*a* → *n, t*) that we have *n* cells after time *t*? – A comprehensive answer to this question is given in ref. [53]. Here, we summarize the results with the addition of more recent results that circumvent issues with computational stability and efficiency.

#### A.1 Exact solutions

For general birth and death rates, *b* ≥ 0 and *d* ≥ 0, and general initial population size *a* ≥ 1, the probability to grow to size *n* in time *t* is given by

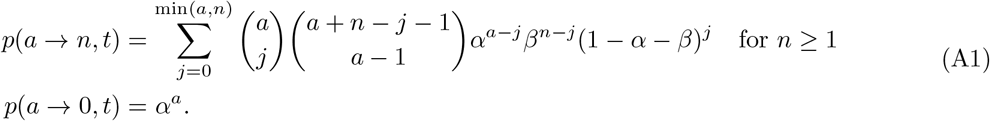

In the super- and subcritical case *b*≠ *d*, we have

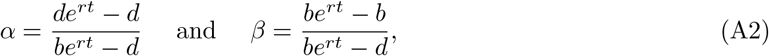

and in the critical case *b* = *d*, we have

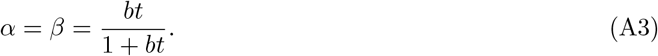

The mean and variance is given by

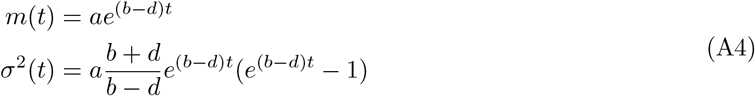

Whereas it is remarkable that an exact solution can be obtained, the solution is computationally expensive and unstable. If the term (1 – *α* − *β*) is negative and the binomial coefficients are large, the limits of machine precision is exceeded and one yields wrong values for *p*(*a* → *n, t*). An alternative form has been derived that sums over positive terms only [12] that reads

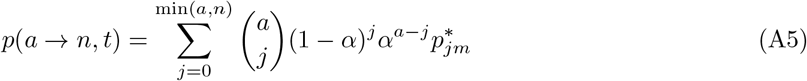

with

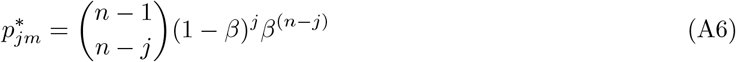

for *j* ≥ 0 and 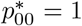 and 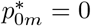 for *j* = 0. Whereas the sum in eqn. (A5) is computationally stable, it is still computationally expensive for large values of *a* or *n*.

If the initial population size is *a* = 1 and *b*≠ *d*, the probability to grow to size *n* after time *t* simplifies to

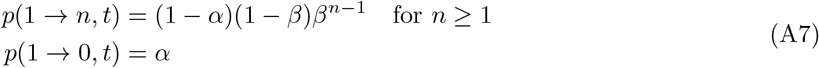

with *α* and *β* given by eqn. (A2). The distribution is recognized as generalized geometric distribution. For the critical case (*b* = *d*) with initial population size *a* = 1, the probability mass becomes

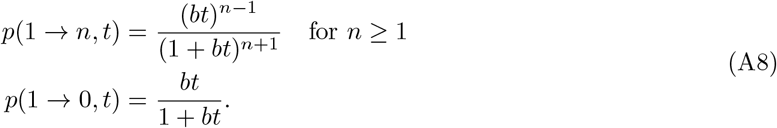

For the pure-birth process (*d* = 0) and general initial population size *a* ≥ 1, we have

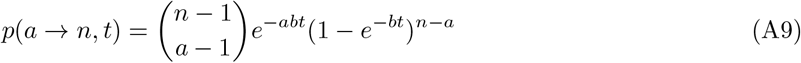

which is a negative binomial distribution, i.e. the probability of having *n* trials given *a* successes with success probability *p* = e^−*bt*^.

For the pure death process *b* = 0, we have

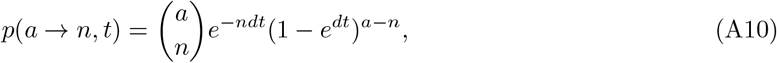

which is a binomial distribution, i.e. the probability of having *n* successes in *a* Bernoulli trials with success probability *p* = e^−*dt*^.

#### A.2 Saddlepoint approximation method

The saddlepoint approximation was introduced to approximate the probability density function or probability mass function given that we know the moment generating function [56] and has found many applications [57].

Consider a discrete random variable *X* that can take values of integers, *k* = 1, 2, 3, Let us denote the moment generating function by *M*(θ), and the cumulant generating function *K*(θ) = log(*M*(θ)). Then the saddlepoint mass function is computed by

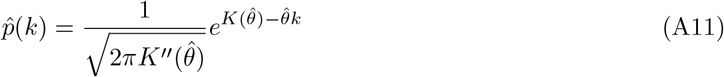

where 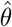 is the solution to 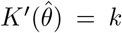. Noteworthy, by design the saddlepoint approximation is not normalized. Once normalized, one talks about the normalized saddlepoint mass function.

Davison et al. applied the saddlepoint approximation on the known moment generating function given by Bailey [13]. For the super- and subcritical case (*b*≠ *d*) the saddlepoint mass function is given by

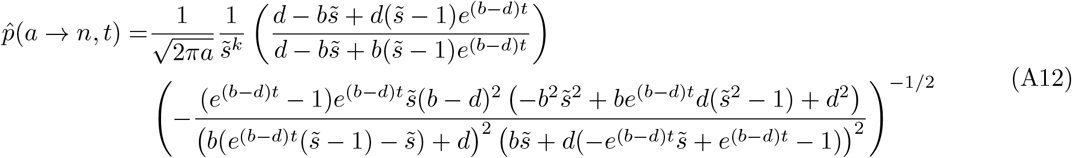

where

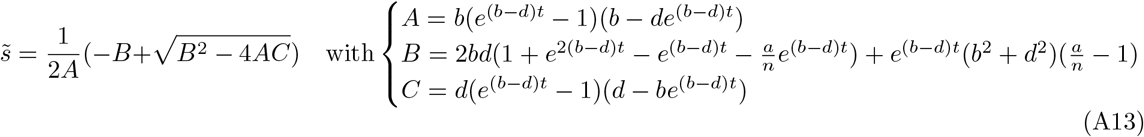

For the critical case (*b* = *d*), it is

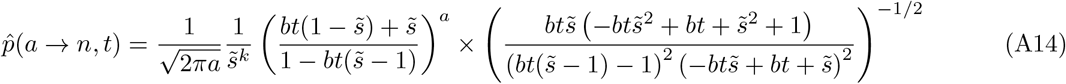

where

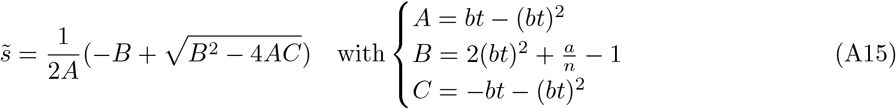

Davison et al. have validated the approximation over a range of parameters. They have found that the approximation yields good results for sufficiently large *n* and *a*. Furthermore, their analysis shows that normalization of the saddlepoint mass function does not lead to significant improvement of the approximation such that we continue working with the unnormalized probabilities.

#### A.3 Heuristic approximation using over-dispersed Poisson distributions

We start by observing that for the pure-birth process, the probability mass follows a binomial distribution. In the pure-death process, it follows a negative binomial distribution which is commonly used to describe overdispersed count data. Furthermore, plotting eqn. (A5) for general *b* and *d* and sufficiently large *n* shows the typical bell shape similar to a Poisson distribution but with increased variance. Those observation motivated us to approximate the probability mass with an overdispersed Poisson distribution that we parameterise with the expressions for the mean and variance.

##### Consul Poisson distribution

We chose the Consul Poisson distribution [58] as one of the simplest formulas of an overdispersed Poisson distribution that reads

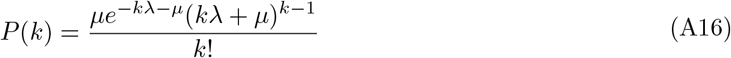

and has mean 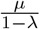 and variance 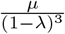. In the case λ = 0, we have the standard Poisson distribution.

We parameterize the Poisson distribution with the mean and the variance of the super- or subcritical birth-death process given in eqn. (A4) that leads to

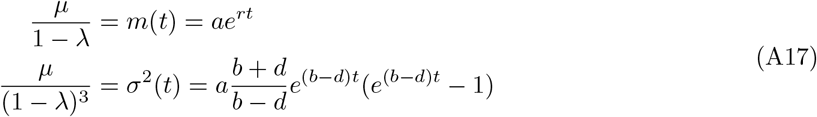

Solving this equation for *µ* and λ has two solutions. We chose the solution leading to positive parameters *µ* and λ in case of overdispersion (i.e. *σ*^2^(*t*) > *m*(*t*)) that is

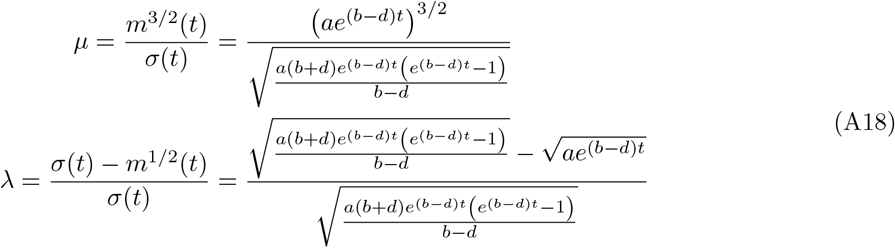

##### Negative binomial distribution

The negative binomial distribution is a standard choice to describe an overdispersed Poisson distribution. It reads

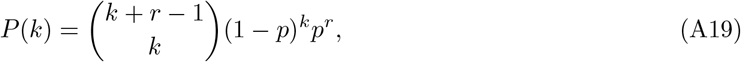

has mean 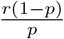 and variance 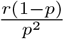. It converges towards a Poisson distribution in the limit *p* → 1. The parameters *p* and *r* can be parameterized and expressed in terms of the mean and variance,

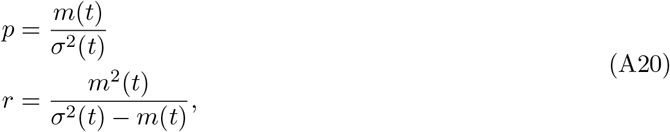

where the mean *m*(*t*) and variance *σ*^2^(*t*) are again taken from eqn. (A4).

### B Probability for the time to reach size *N*

The probability for the time to reach size *N* starting from one cell (*a* = 1) was derived in ref. [15]. The probability density is approximately

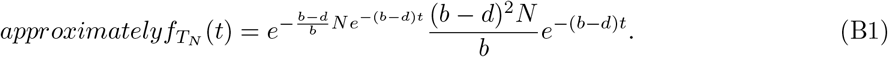

This is a Gumbel distribution, which is commonly written in terms of a location parameter *µ* and shape parameter *β*. We can write the density as

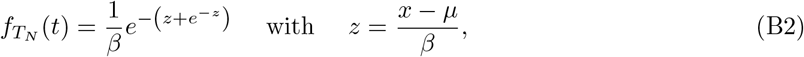

such that 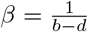 and 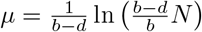.

We can then make use of the known properties of the Gumbel distribution. In particular, the expectation is

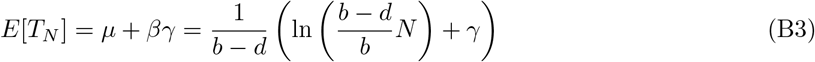

where *γ* ≈ 0.577 is the Euler constant. Further, the moment-generating function defined by

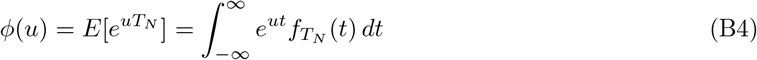

takes the form

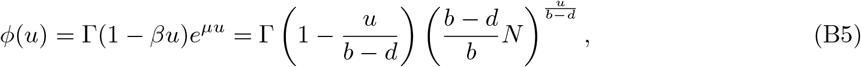

Where 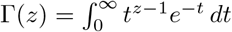 is the gamma function.

The Gumbel distribution is defined over (−∞, ∞) whereas we are only interested in positive times *T*_*N*_ > 0. However, in applications the integral over the density for negative *t* is typically negligible and we proceed with approximating 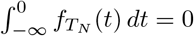.

### C Switching between fixed-time and fixed-size conditioning

The random process is stopped at fixed time *T* = *t* or fixed size *N* = *n* and an observable is measured. We know that the probability to grow to size *n* in time *t* (Appendix A) that we write as *p*(*n*|*t*) to be explicit on the conditioning. Similarly, starting from a single cell, we know the probability for the time *t* to grow to size *n* for the first time (Appendix B) that we denote by f_*T*_ (*t*|*N* = *n*). We investigate the expectation of an observable *O*.

Given we know the expected observation for fixed time *T* = *t*, we can compute the expected observation for fixed size *N* = *n* by

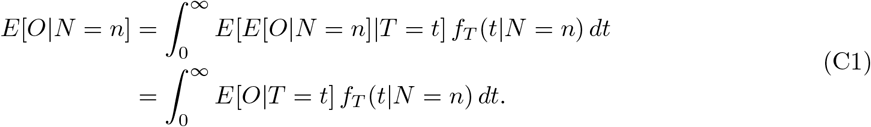

In the first equality, we decompose the expectation of *E*[*O*|*N* = *n*] into expectations for all possible times *t*. In the second equality, we use the law of total expectation which states *E*[*E*[*O*|*N* = *n*]|*T* = *t*] = *E*[*O*|*T* = *t*].

Similarly, if we know the expected observation for fixed size *N* = *n*, we can compute the expected observation for fixed time *T* = *t* by

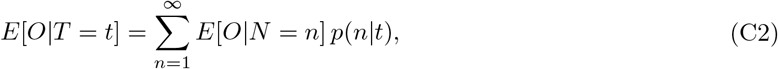

where we condition on survival and thus chose *n* = 1 rather than *n* = 0.

### D Proof of theorem 1

**Part 1:** We prove eqn. (6) for general *N*_0_ following Appendix B of ref. [16] who proved the result for *N*_0_ = 1. Restricted (but not conditioned) on survival of the entire population {Z_0_(*t*) > 0}, the expected number of mutations that emerged in the time interval [*t*^′^, *t*^′^ + *dt*^′^] and grow to size *k* in the remaining time *t* − *t*^′^ was shown to be

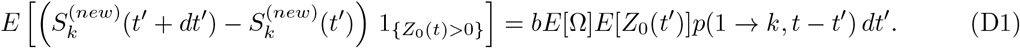

Here, Ω is the number of mutations generated in a division. Given our *Theoretical framework*, we have *E*[Ω] = 2*m*. The expected population size in the birth-death process is given by *E*[Z_0_(*t*^′^)] = *N*(*t*^′^) = *N*_0_*e*^(*b*−*d*)*t*^*′*. Conditioning the total population to survive time *t* leads to

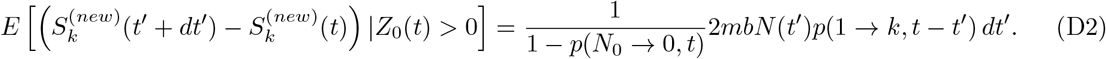

It remains to integrate over the time interval of entire time interval *t*^′^ ∈ [0, *t*] such that

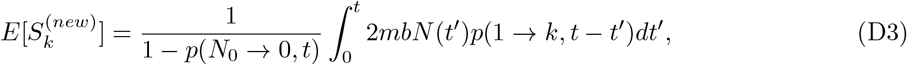

which makes the proof complete.

Noteworthy, the expected BWD is given by

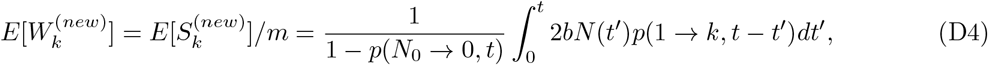

**Part 2:** Next, we prove eqn. (7). We start by writing the initial BWD as 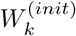, label branches of the phylogenetic tree with *i* = 1, 2, …*L* and their branch widths with *k*_*i*_. During time *t*, branches grow stochastically and we filter for those that grow to size *k* by writing

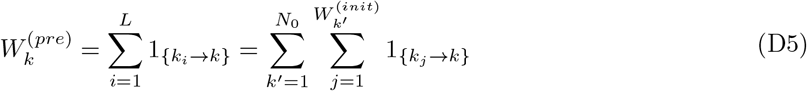

where 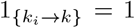 if branch *i* of width *k*_*i*_ grows to size *k* and 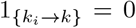 otherwise. In the second equality, we used that the *L* branches are partitioned into *N*_0_ classes 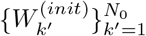 according to their initial width *k*^′^.

Taking the expectation, we have

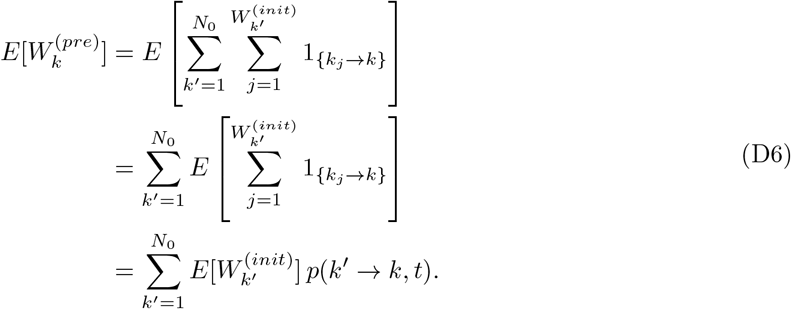

In the second equality, we used linearity of the expectation. In the third equality, we used Wald’s identity together with 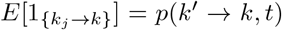 when *k*_*j*_ has width *k*^′^. We substitute 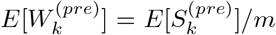 and 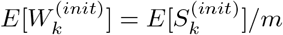 and condition the entire population on survival such that

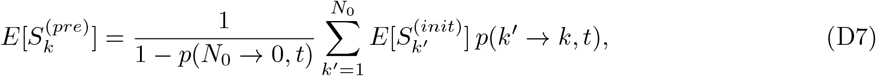

which is the desired result.

### E Site frequency spectra at detection

#### E.1 Fixed-time expectation

Starting from a single cell with *u*_0_ mutations, the population grows time *t*_*d*_ to detection. The average growth in the time interval [0, *t*_*d*_] is given by 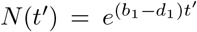 and the expected population size at detection conditioned on survival is 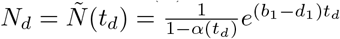.

##### Newly emerging mutations

Following theorem 1, we have

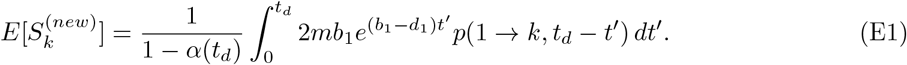

In ref. [17] and [16], the integral is rewritten and simplified for the limits *N*_*d*_ → ∞ and *k* → ∞ giving us

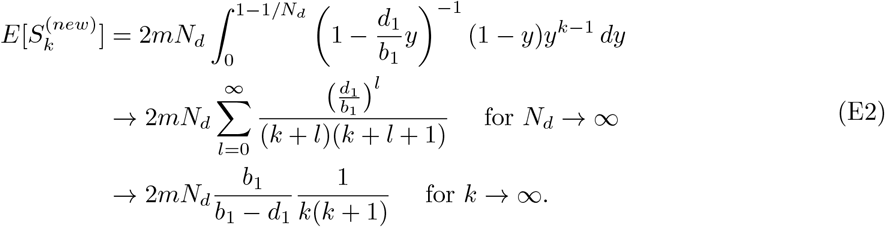

Considering only mutations with infinite lineage, one can use the probability mass functions of the pure-birth process and the SFS was shown to be [16]

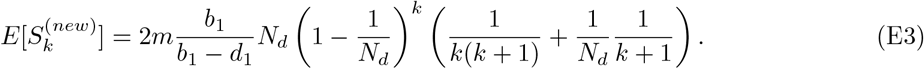

Since all mutations have an infinite lineage in the pure-birth process, this formula also provides the SFS for the pure-birth process when setting *d*_1_ = 0.

##### Preexisting mutations

Following theorem 1, we have

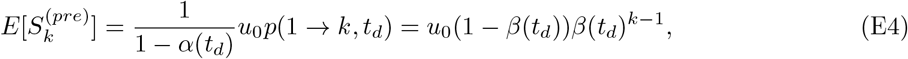

which is just the number of initial mutations multiplied by the probability of growing from size 1 to size *k* in time *t*_*d*_ conditioned on survival. *β*(*t*_*d*_) is defined in eqn. (A2).

Our main interest is in mutation accumulation during tumour progression. Therefore, we work with *u*_0_ = 0 in the main text leading to 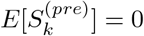 and 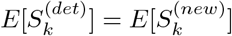.

#### E.2 Fixed-size expectation

Following eqn. (C1), the expected SFS at fixed size 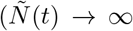 can be expressed in terms of the expected SFS at fixed time *T* = *t* by writing

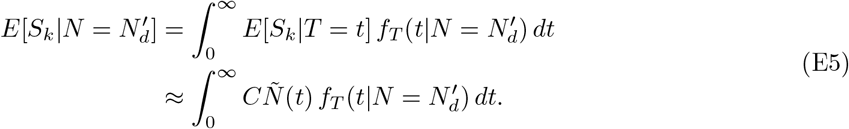

In the second line, we used the solution for large population size (*Ñ*(*t*) → ∞ in eqn. (8)) such that *E*[*S*_*k*_|*T* = *t*] is proportional to the expected size at time *t*. Here, C is independent of time and reads

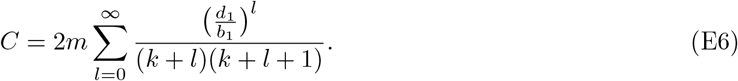

The expected size at fixed time *t*_*d*_ can be written as

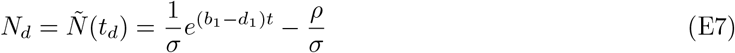

With 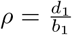 and 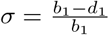 (eqn. 17 in ref. [16]). The integral can be expressed in terms of the moment generating function (eqn. (B5)) such that

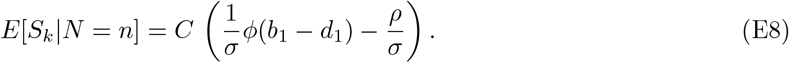

However, we have

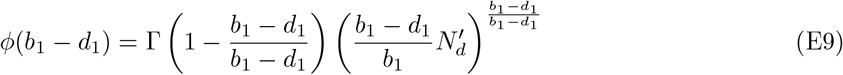

and Γ(0) is not defined.

To avoid the convergence problem, we consider the normalized SFS that we define by 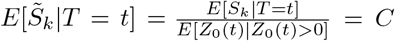. Again, we switch from the fixed-time expectation to the fixed-size expectation, which yields

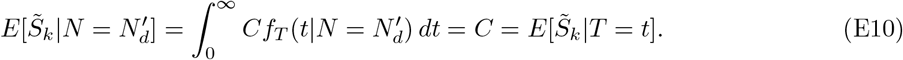

We find that the fixed-time expectation and fixed-size solution expectation. The perfect one-to-one correspondence vanishes if we take the exact expression for the fixed-time expectation instead of the large-size approximation in eqn. (8).

### F Site frequency spectrum after treatment with homogeneous response

We compute the expected SFS after treatment with homogeneous treatment response. As initial condition, we consider fixed size 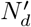 and the expected SFS at detection. For the initlal SFS, we approximate the fixed-size expectation of the SFS with the fixed-time expectation of the SFS by replacing the expected size *N*_*d*_ = *Ñ*(*t*_*d*_) with the fixed-size 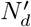 in eqn. (8). We denote the treatment time by *t*_*f*_ after which the population conditioned on survival has expected size 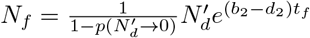.

#### F.1 Decreasing cell populations

We consider decreasing cell populations characterized by *b*_2_ < *d*_2_ and 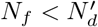.

##### Newly emerging mutations

Following theorem 1, we have

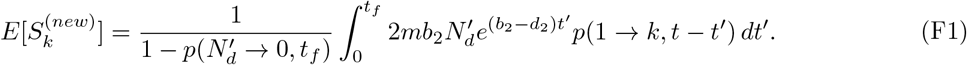

Here, *p*(1 → *k, t* − *t*^′^) is given by eqn. (A7). Using the same transformation as for the exponential growth starting from a single cell (ref. [16] eqn. (B.5) to (B.6)), the integral can be rewritten as

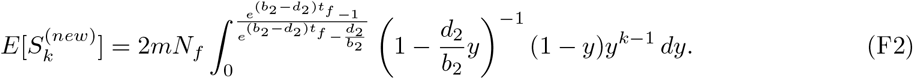

In general, the upper integral bound does not go to 1 such that we cannot transform the integral into a sum as done in ref. [16]. Instead, we solved the integral numerically.

The situation changes in the limit *b*_2_ → *d*_2_. Then, the upper integral bound becomes 1 but at the same time the integrand becomes *y*^*k*−1^ resulting in 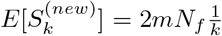. This result coincides with the equilibrium solution for *b*_2_ = *d*_2_ that will be further refined in the next subsection (see eqn. (F5)).

In the limit 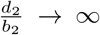, the upper integral bound converges to 0 whereas the integrand converges to 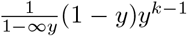 resulting in 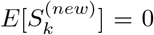. The result is intuitive. If there are no birth events, then there are no new mutations.

##### Preexisting mutations

Following theorem 1, we have

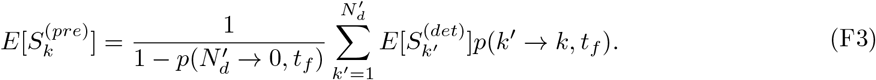

Here, *p*(*k*^′^ → *k, t*_*f*_) is given by eqn. (A1) parameterized with *b*_2_ and *d*_2_. By construction, the initial SFS is the SFS at detection whose expectation is given by eqn. (8).

In general, the exact probabilities *p*(*k*^′^ → *k, t*_*f*_) take a complicated form. Consequently, the evaluation of 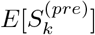 for specific *k* consists of computing a sum ranging from 0 to 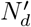 over sums ranging from 0 to min(*k, k*^′^). The computational cost is high but can be decreased with the approximation methods discussed in Appendix A.

In the pure-death process with *b*_1_ = 0, we can use (A10) that resolves the second sum without the use of approximation methods. Resolving the first sum remains complicated but is possible when we condition on fixed size instead of fixed time (see Appendix G).

#### F.2 Constant cell populations

Next, we consider cell populations that remain approximately constant characterized by *b*_2_ = *d*_2_ and 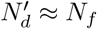.

##### Newly emerging mutations

Following theorem 1, we have

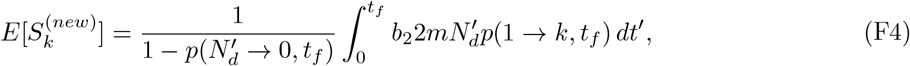

where *p*(1 → *k, t*_*f*_) are given by eqn. (A8). We solve the integral and obtain

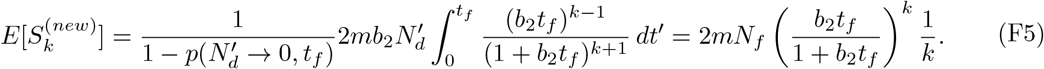

As *t* → ∞, the SFS is proportional to 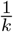, which is known as the equilibrium solution from other constant population size models, namely the Moran process or the Wright-Fisher process [20].

##### Preexisting mutations

Following theorem 1, we have

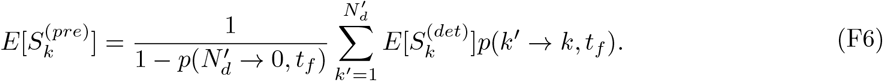

Here, *p*(*k*^′^ → *k, t*_*f*_) is given by eqn. (A1) parameterized with *b*_2_. Again, 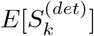 is given by eqn. (8).

Similar to the decreasing population, the computation of 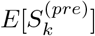 for one specific *k* consists of evaluating the sum of sums where the second sum can be resolved using the approximations described in Appendix A.

#### F.3 Increasing cell populations

Eventually, we consider increasing cell populations characterized by *b*_2_ > *d*_2_ and 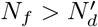.

##### Newly emerging mutations

The exact expressions for increasing populations coincide with the ones for decreasing populations such that

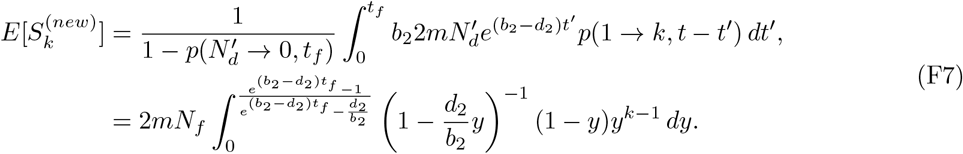

We solved the integral numerically.

In the limit *t*_*f*_ → ∞, the upper integral bound converges to 1 and following ref. [16] (computations after eqn. (B.6)), we can express the integral as infinite sum

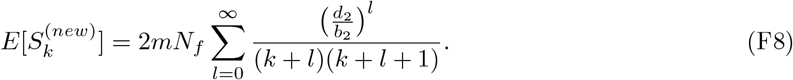

This limit has restricted application. After detection, the cancer is unlikely to continue growing over a long time since the cancer likely becomes lethal by then. However, this scenario is applicable in case birth and death rates are changed before detection, e.g. through activation of the immune system, angiogenesis or other environmental factors.

In the limit *b*_2_ → *d*_2_, we recover the 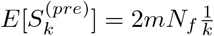 like we did for decreasing populations.

In the limit *d*_2_ → 0, we have

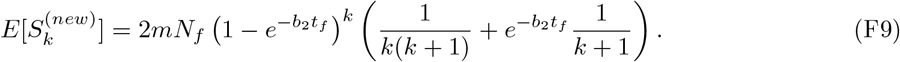

##### Preexisting mutations

Again, the expressions for increasing populations coincide with the ones for decreasing populations such that

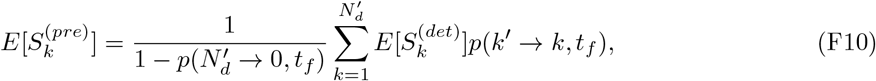

where *p*(*k*^′^ → *k, t*_*f*_) is given by eqn. (A1) parameterized with *b*_2_ and *d*_2_, and 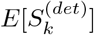 is given by eqn. (8). Obtaining a single value comes with evaluation of two sums where the second sum can be resolved using approximation methods for *p*(*k*^′^ → *k, t*_*f*_) in Appendix A.

#### F.4 Approximating the fixed-size expectation with the fixed-time expectation

For increasing or decreasing populations, we approximate the expected SFS at fixed size 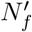 with the expected SFS at fixed time *t*_*f*_. Therefore, we transform the treatment time into size after treatment using the expression for the expected size after treatment for time *t*_*f*_ giving us

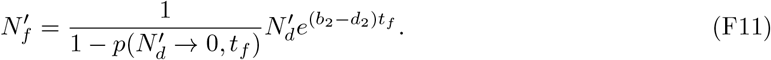

Inverting the exact expression analytically is complicated. However, in applications we have 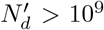 and given that we still have an observable cancer *N*_*f*_ > 10^3^. For population sizes of this magnitude, we have 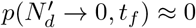 such that 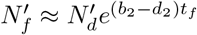 and

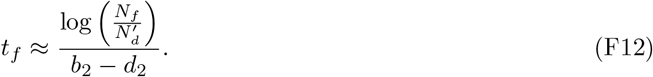

We inserted this expression into the fixed-time expectation to approximate the fixed-size expectation.

### G The site frequency spectrum retains its *k*^−2^ tail after treatment with homogeneous response

After treatment of cancers with homogeneous treatment response, the expected SFS retains its power law tail *E*[S_*k*_] ∼ *k*^−2^. We assume that mutations that occur in less than 1% of the population are not observable due to sequencing limitations. Thus, we define the tail distribution by S_*k*_ with *k* ≥ *k*_min_ = 1%*N*. First, we show that the *k*^−2^ tail in retained in simulated data that have population sizes in the magnitudes 10^4^ − 10^5^. Second, we make a parameter estimation for realistically large tumors and argue why the *k*^−2^ tail in retained. We show that newly emerging mutations remain at low unobservable frequencies, and observable preexisting mutations are usually large and behave nearly deterministic.

#### G.1 Observation of the *k*^*−*2^ tail in simulated data

We observe a power law of the form *E*[S_*k*_] ∼ *k*^−*γ*^ from fig. 2D-F by noting of the plots build a straight line in the loglog-plot for sufficiently large *k*. Next, we infer the exponent *γ* from the simulated data using the maximum likelihood estimator from ref. [55] (see *Methods*). We first infer exponents of simulated data at detection (fig. S1A-D). As expected, we obtain exponents *γ* ≈ 2. Depending on the cutoff *k*_min_, we observe a small bias towards larger exponents (fig. S1B and D). This bias may be explained by the high variability in the largest site frequencies. We also note that strictly speaking, we did not show that the SFS of a single realization follows a power law distribution but the expected SFS does.

Next, we inferred the exponents in simulated data after treatment with homogeneous response. Again, we find exponents *γ* ≈ 2 with small biases towards larger exponents consistent with the inference artefact that we also obtained for the SFS at detection. We conclude the observation of the S_*k*_ ∼ *k*^−2^ tail after treatment with homogeneous response.

#### G.2 Estimating biologically reasonable parameters

We assume that a cancer is detected at population sizes in the range *N*_*d*_ = 10^9^ − 10^12^. Assuming a typical division time τ_2_ = 5 days, the birth rate can be estimated to be 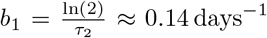. For cancers growing to detection size in around 5-15 years, the growth rate is in the range from 0.005 − 0.015 days^−1^ translating into death rates close to the birth rate in the range *d*_2_ = 0.125 − 0.135 days^−1^. In the following, we work with an intermediate death rate *d*_1_ = 0.13 days^−1^.

We set treatment time to *t*_*f*_ = 180 days. We assume that treatment affects the death rate only such that we keep the birth rate *b*_2_ = 0.14 days^−1^. The death rate is adapted according to the three treatment scenarios. In an effective treatment, the cancer may decrease to 1% of its original size translating into death rate *d*_2_ ≈ 0.166 days^−1^. For approximately constant disease, we have *d*_2_ = 0.14 days^−1^.

For ineffective treatment, we consider the case in which treatment has no effect at all such that *d*_2_ = 0.13 days^−1^.

#### G.3 Decreasing cell populations

##### In the pure-death process, the expected SFS has a *k*^−2^ tail

If *b*_2_ = 0.0, then there are no newly emerging mutations such that 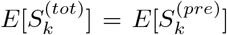. Instead of conditioning the expectation on fixed time *t*_*f*_, let us condition on fixed size 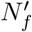 starting from fixed detection size 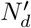. In the pure-death process, growing from size *k*^′^ to size *k* during treatment is the same as sampling *k* cells from *k*^′^ cells without replacement. Thus, the probability to grow to size *k* is given by the hypergeometric distribution,

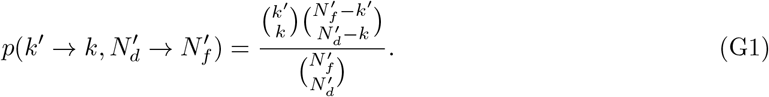

If 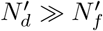, the hypergeometric distribution can be approximated by the binomial distribution (sampling with replacement), which is

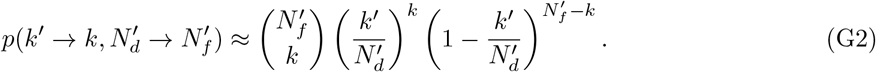

Interestingly, we arrive at a binomial distribution like the probability mass function for fixed-time in eqn. (A10), although the binomial distribution is different parameterized.

From here, we can follow the derivation of Durrett who studied the form of the sampled SFS to describe frequencies in a sample (theorem 3 in ref. [15]). We transform the sum for the SFS into an integral and focus on the tail of the SFS at detection such that

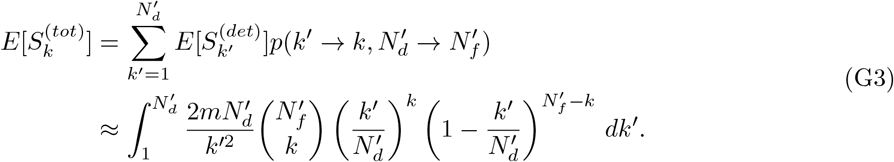

We substitute the initial mutant size with its frequency, 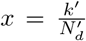 and 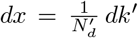. Furthermore, we assume 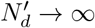 such that the lower integral boundary goes to 0. We have

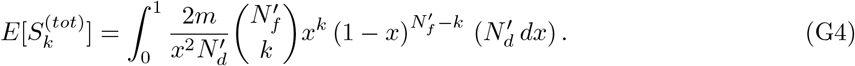

Using that 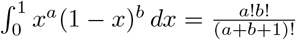 (see Lemma 2 in ref. [15]), we obtain

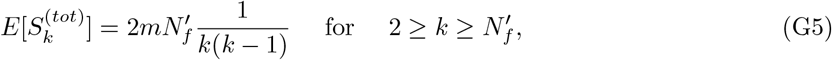

which scales as *k*^−2^ for large *k*. Noteworthy, the continuity approximation works only for sufficiently large *k*.

##### The SFS of preexisting mutations retains its *k*^−2^ tail

Consider a cancer cell population with detection size *N*_*d*_ = 10^12^ that after treatment for time *t*_*f*_ = 180 days shrinks to 1% of its original size, *N*_*f*_ = 10^10^. We compute the mean and the standard deviation for a single site frequency that is initially in 1% of the cells, i.e. *k*^′^ = 10^10^. Inserting the birth and death rates from our parameter estimation *b*_2_ = 0.14 days^−1^ and *d*_2_ = 0.166 days^−1^ into eqn. (A4), we have

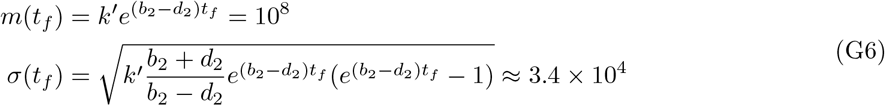

Starting from a large populations, we know that the probability mass function takes a bell-shaped curve as long as we are not too close to the extinction boundary [13]. Observing that the standard deviation is multiple magnitudes lower than the mean, we conclude a sharply peaked probability mass function that we interpret as approximately deterministic behaviour.

If the growth dynamics of site frequencies is deterministic, it is straightforward to show that the initial shape of the SFS is retained. Focusing on the tail of the SFS at detection and assuming deterministic growth during treatment, we have

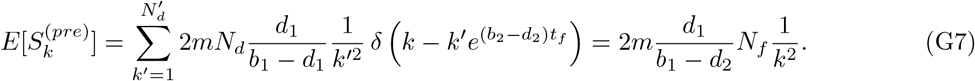

Here, we use deterministic growth for the total population 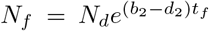 and approximate the probability mass function by 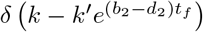 that is 1 if 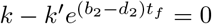 and 0 otherwise.

##### Newly emerging mutations are restricted to small frequencies

Since *b*_2_ < *d*_2_, newly emerging mutations survive only due to drift. Given *b*_2_ = 0.14 and *d*_2_ ≈ 0.166, the probability to survive time *t*_*f*_ is only 1 − *α*(*t*_*f*_) ≈ 0.000243. Further, the probability to reach larger sizes *k* > 1 is decreasing with time.

#### G.4 Constant cell populations

##### The SFS of preexisting mutations retains its *k*^−2^ tail

Consider a cancer cell population with detection size *N*_*d*_ = 10^12^ that remains approximately constant over treatment time *t*_*f*_ = 180 days. We compute the mean and the standard deviation for a single site frequency that is initially in 1% of the cells, i.e. *k*^′^ = 10^10^. Inserting the birth and death rates from our parameter estimation *b*_2_ = *d*_2_ = 0.14 days^−1^ into eqn. (A4), we have

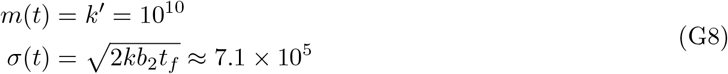

With analogous argumentation to the decreasing population, we conclude that the growth is approximately deterministic and the SFS of preexisting mutations retains its *k*^−2^ tail.

##### Newly emerging mutations are restricted to small frequencies

Using the same parameters as before, we compute the probability that a newly emerging mutation grows to abundance *k* > 100 during treatment time *t*_*f*_ = 180 days

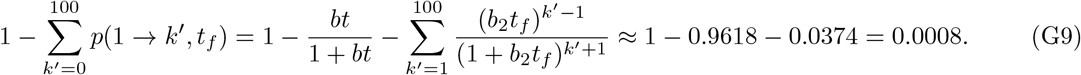

The chance is very small, and we conclude that genetic drift alone is not enough to grow to observable site frequencies in the order *k* = 10^10^.

This is also reflected in the expression 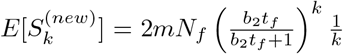, which has an exponential cutoff with *k*. Inserting the parameters above, we note that there is a dominating exponential cutoff with *k* that reads

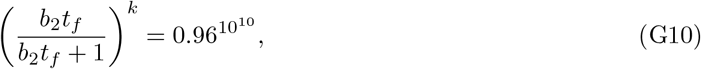

which is extremely small such that 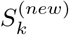 is virtually zero for large *k*.

#### G.5 Increasing cell populations

##### The SFS of preexisting mutations retains its *k*^−2^ tail

Consider a cancer cell population with detection size *N*_*d*_ = 10^11^ that after ineffective treatment for time *t*_*f*_ = 180 days continues increasing to approximately *N*_*f*_ *≈* 6 × 10^11^. We compute the mean and the standard deviation for a single site frequency that is initially in 1% of the cells, i.e. *k*^′^ = 10^9^. Inserting the birth and death rates from our parameter estimation *b*_2_ = 0.14 days^−1^ and *d*_2_ = 0.13 days^−1^ into eqn. (A4), we have

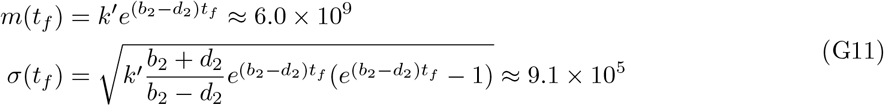

With analogous argumentation to the decreasing population, we conclude that the growth is approximately deterministic and the SFS of preexisting mutations retains its *k*^−2^ tail.

##### Newly emerging mutations are restricted to small frequencies

In contrast to decreasing and the constant populations, newly emerging mutations grow in their abundance not only to due to genetic drift. However, since the cancer is usually detected at very large sizes, and preexisting site frequencies grow at the same rate, it is highly unlikely that newly emerging mutations grow to 1% of the population. With the estimated birth and death rate, we have mean and standard deviation for the abundance of a newly emerging mutation growing for time *t*_*f*_ = 180 days,

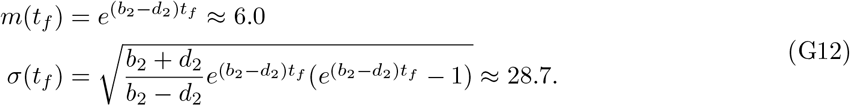

Although the standard deviation is very large compared to the mean, the likelihood to reach observable size *k*_min_ ≈ 6.0 × 10^9^ is again virtually zero.

### H Proof of theorem 2

**Part 1:** We prove eqn. (15) building up on the proof of theorem 1. Restricted (but not conditioned) on survival of the entire population {*Z*_0_(*t*) > 0}, the expected number of mutations that emerged in the time interval [*t*^′^, *t*^′^ + *dt*^′^] and survive in the remaining time *t* − *t*^′^ is

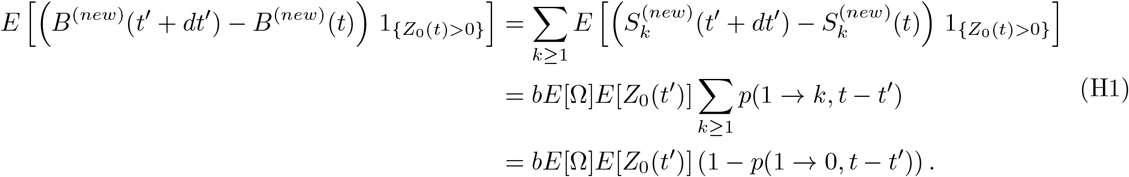

In the first equality, we used that 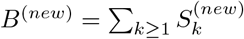 together with linearity of the expectation. The second equality is equivalent to the first equation in the proof of theorem 1 taken from ref. [16]. Again, Ω is the number of mutations generated in a division such that *E*[Ω] = 2*m* and the expected population size is 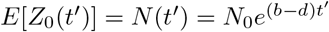. After conditioning the total population to survive time *t*, we have

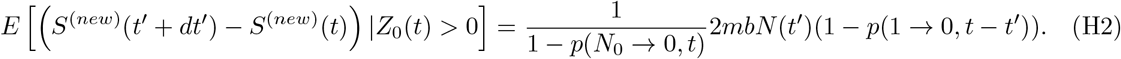

We integrate over the time interval of interest *t*^′^ ∈ [0, *t*] giving us

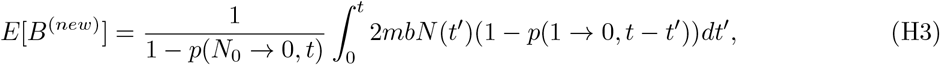

which makes the proof complete.

The expected number of divisions is

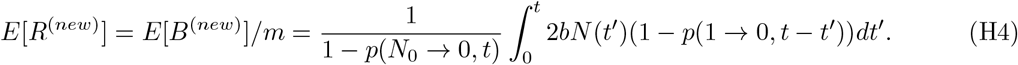

**Part 2:** Next, we prove eqn. (16). Analogously to the proof of the second part of theorem 1, we start by writing the initial BWD as 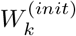, label its *L* branches with index *i* = 1, 2, …*L* and their branch widths with *k*_*i*_. During time *t*, branches may go extinct and we count branches leading to at least one living cell that is

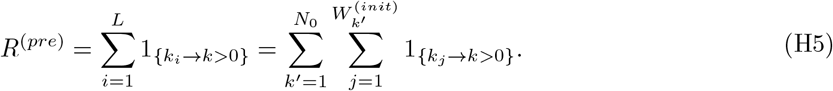

Here, 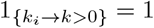 if branch *i* of initial width *k*_*i*_ grows to size *k* > 0 and 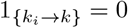 otherwise. Again, we used that the L branches are partitioned into *N*_0_ classes 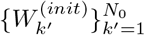 according to their initial width *k*^′^.

Taking the expectation and using the same steps as in the proof of theorem 1, we have

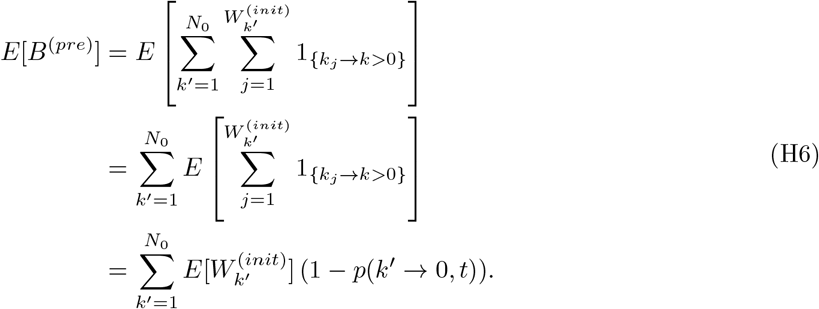

We substitute *E*[*R*^(*pre*)^] = *E*[*B*^(*pre*)^]/*m* and 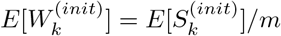 and condition the entire population on survival such that

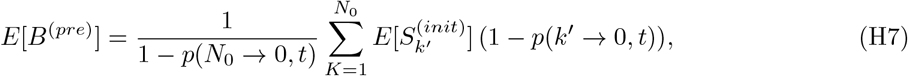

which is the desired result.

### I The total mutational burden at detection

#### Pure-birth process

If we consider the fixed-size limit, we know that there are exactly *N*_*d*_ − 1 divisions that a population must undergo to grow form 1 cell to *N*_*d*_ cells. Thus, the expected tMB is

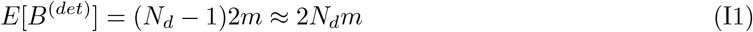

Furthermore, since each division comes with *µ*_*i*_ ∼ Poiss(*m*) mutations, we know that the tMB is Poisson distributed

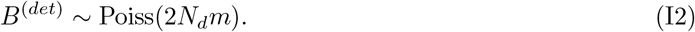

If there were *u*_0_ mutations in the ancestor cell, one simply needs to add them as constant outside the Poisson distribution, *B*^(*det*)^ ∼ Poiss(2*N*_*d*_*m*) + *u*_0_.

In the fixed-time condition, we consider a population growing time *t*_*d*_ at which we expect population size *N*_*d*_ = *E*[*Z*_0_(*t*_*d*_)]. This translates into an expected number of *N*_*d*_(*t*_*d*_) − 1 divisions similar to the fixed-size expectation. However, the distribution of *B*^(*det*)^ is not Poisson distributed anymore. The number of divisions differs between realizations thus leading to additional stochastic effect.

##### Birth-death process

Following theorem 2, starting from a single cell with no mutations that grows for time *t*_*d*_ and is conditioned to survive, we have

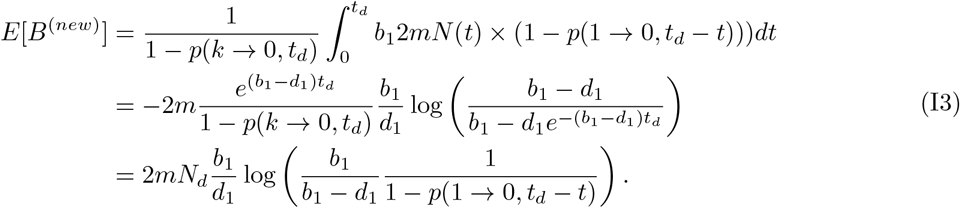

We can further manipulate the expression to show agreement with ref. [16]. We have

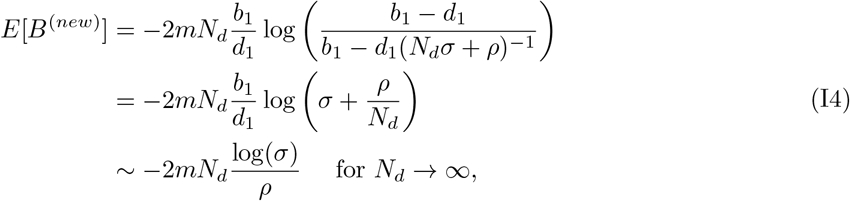

where we used that 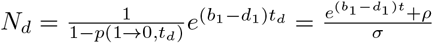.

If there be *u*_0_ mutations preexisting in the first cell, they simply need to be added as a constant such that total tMB is *E*[*B*^(*tot*)^] = *E*[*B*^(*new*)^] + *u*_0_.

### J The total mutational burden after treatment with homogeneous response

We compute the expected tMB after treatment with homogeneous treatment response. As initial condition, we consider fixed size 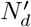 and the expected SFS at detection. For the initial SFS, we approximate the fixed-size expectation of the SFS with the fixed-time expectation of the SFS by replacing the expected size *N*_*d*_ = *Ñ*(*t*_*d*_) with the fixed-size 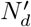 in eqn. (8). We denote the treatment time by *t*_*f*_ after which the population conditioned on survival has expected size 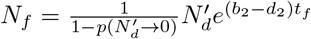. Since the extinction probability appears in the final expressions, it is convenient to write *α*(*t*) for *p*(1 → 0, *t*), and *α*^*a*^(*t*) for *p*(*a* → 0, *t*).

#### J.1 Decreasing cell populations

We consider decreasing cell populations characterized by *b*_2_ < *d*_2_ and 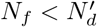.

##### Newly emerging mutations

Following theorem 2, we have

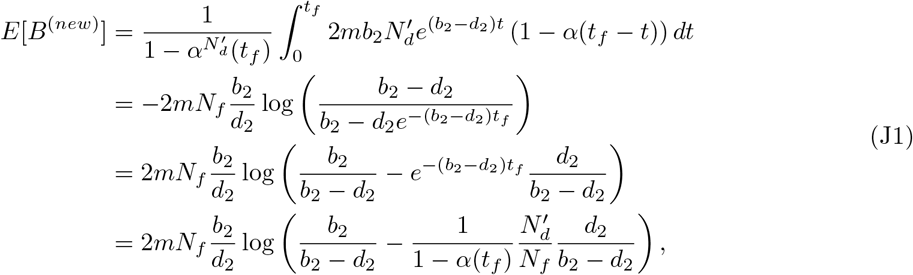

For *t*_*f*_ → ∞, the expression coincides with the the tMB at detection. In particular, it is independent of the initial population size 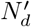. This makes sense. Newly emerging mutations are only short-lived and the major contributions to *B*^(*new*)^ are very recent mutations.

##### Preexisting mutations

Following theorem 2, we have

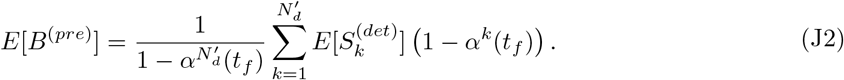

Here, *α*(*t*) is given by eqn. (1) parameterized with *b*_2_ and *d*_2_. By construction, the initial SFS is the SFS at detection whose expectation is given by eqn. (8).

#### J.2 Constant cell populations

Next, we consider cell populations that remain approximately constant characterized by *b*_2_ = *d*_2_ and 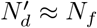.

##### Newly emerging mutations

Following theorem 2, we have

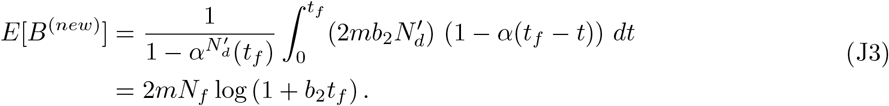

##### Preexisting mutations

Following theorem 2, we have

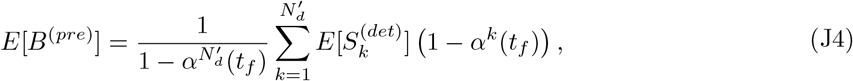

Again, *α*(*t*) is given by eqn. (1) parameterized with *b*_2_ and *d*_2_. The initial SFS is given by eqn. (8).

#### J.3 Increasing cell populations

Eventually, we consider increasing cell populations characterized by *b*_2_ > *d*_2_ and 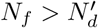.

##### Newly emerging mutations

The expressions for increasing populations coincide with the ones for decreasing populations such that

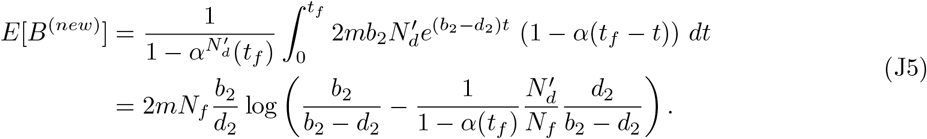

In the limit 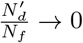, the expression is independent

##### Preexisting mutations

Following theorem 2, we have

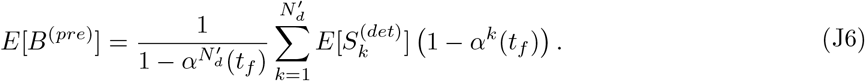

Once more, *α*(*t*) is given by eqn. (1) parameterized with *b*_2_ and *d*_2_. The initial SFS is given by eqn. (8).

#### J.4 Approximation for preexisting mutations

We resolve the sum for *E*[*B*^(*pre*)^] using diverse approximation. In this section, the extinction probability will always be the extinction probability after time *t*_*f*_ such that we write *α* = *α*(*t*_*f*_) that are parameterized with *b*_2_ and *d*_2_.

##### Consider 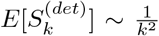 as expected SFS at detection

For large site frequencies *k*, the SFS at detection is approximately 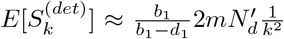. Using this expression for *E*[*B*^*pre*^] and with help of *Wolfram Mathematica*, we obtain

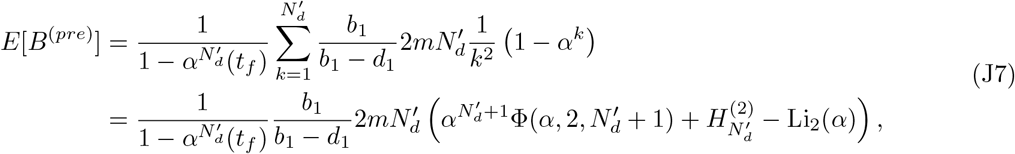

where 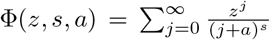 is the Lerch phi, 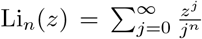 is the polylogarithm and 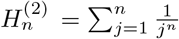 are generalized harmonic numbers. For sufficiently large 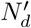, the first term in the parentheses becomes very small and we are left with

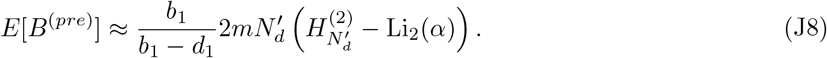

Notably, the 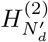 is time-independent and positive whereas −Li_2_ (*α*) is time-dependent and negative. We can interpret the first term as the number of mutations at detection which we can also obtain directly by summing over 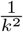. Then, the second term describes the number of mutations that are lost over time.

##### Consider 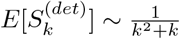 as expected SFS at detection

Most mutations occur at low abundance where 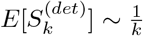 is not valid. To get one step closer to the exact expected SFS at detection, we next consider 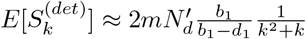. Using *Wolfram Mathematica*, we obtain

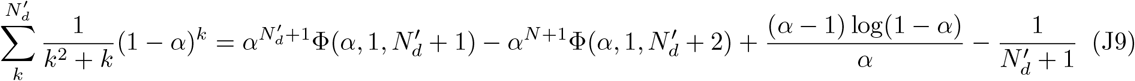

For large 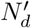, the first two terms and the last term become very small leaving us with

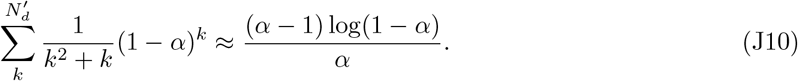

Using this expression to compute the tMB of preexisting mutations, we obtain

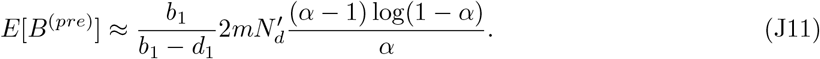

##### Heuristic adjustment at *t*_*f*_ = 0

At *t*_*f*_ = 0, the extinction probability is *α* = 0, and

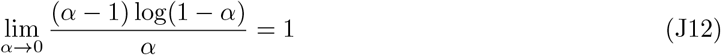

such that 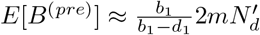 at time *t*_*f*_ = 0. This is not accurate for the general birth-death process as we know from the section on the tMB at detection. Requiring a correct result at time *t*_*f*_ = 0, we make a heuristic correction and write

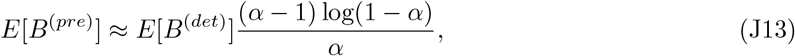

where *E*[*B*^(*det*)^] is given by eqn. (17).

### K Proof of theorem 3

**Part 1:** We prove eqn. (22). Taking the expectation of the scMB described by eqn. (21), we have

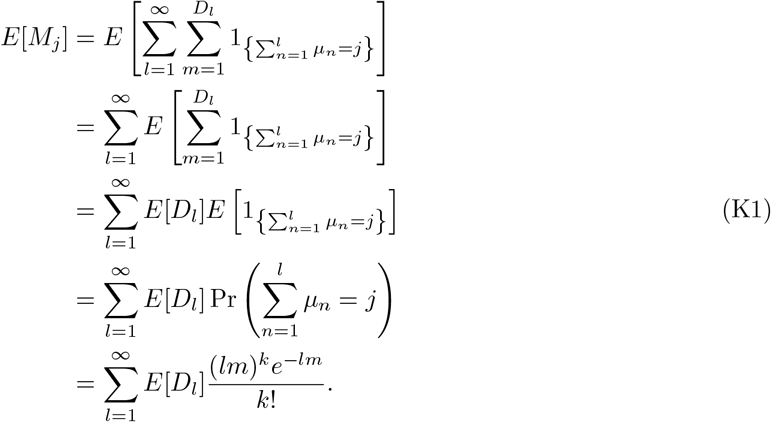

**Part 2:** Next, we prove eqn. (23). We denote the number of cells with *l* divisions at time *t* by *D*_*l*_(*t*). We follow the derivation of ref. [5]. We start from a population with *N*_0_ cells at time *t* = 0 such that *D*_0_(*t* = 0) = *N*_0_. Considering only the mean behaviour, we write the following differential equations for *D*_*l*_

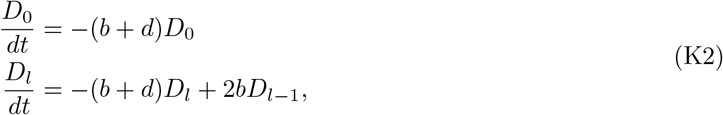

which has solutions

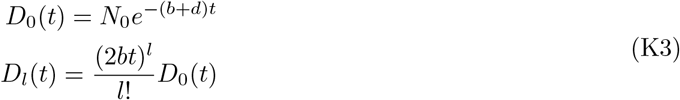

The expected population size after time *t* is *N* = *N*_0_*e*^(*b*−*d*)*t*^, which leads to the desired result.

### L Average number of divisions in the population

#### L.1 Increase of the average generation per division

The following derivation was motivated by ref. [9]. Consider a population of *N* cells, where the number of divisions that each cell has undergone are saved in a vector 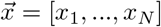 with mean 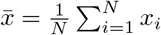. After one division, the population has increased to size N +1 and we have division vector 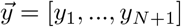 with mean 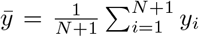. Let us denote the index of the dividing cell by *j* such that *x*_*i*_ = *y*_*i*_ for *i* ≠ *j* and *y*_*j*_ = *y*_*N*+1_ = *x*_*j*_ + 1. Then, we have

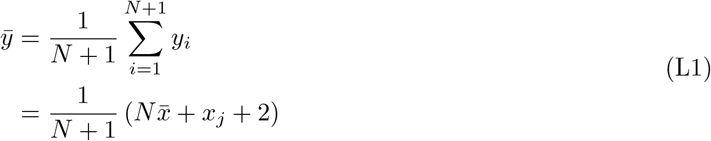

If we now average over many realizations, then

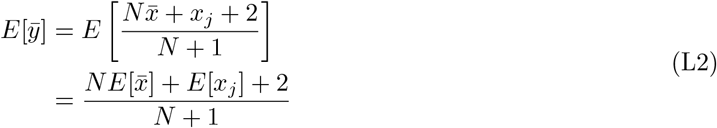

Now, note that

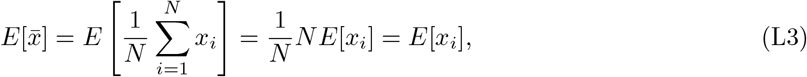

such that we have

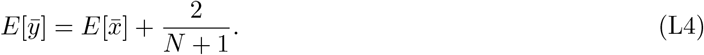

Thus, the average increase for going from size *N* to size *N* + 1 is given by 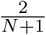.

#### L.2 Fixed-size solution in the pure-birth process

For a pure-birth process starting from 1 cell with 0 divisions, we have *N* − 1 divisions to grow to size *N* where the *i*-th division increases the generation average by 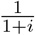 such that the expected average number of division reads

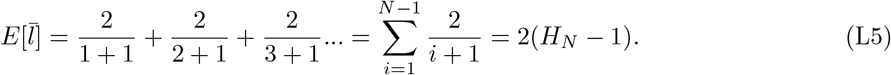

More generally starting from *N*_0_ cells with 0 divisions growing to size *N*, we have

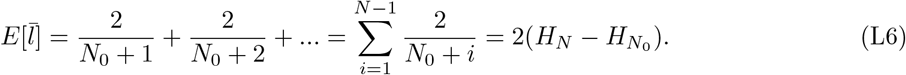

#### L.3 Fixed-time solution in the pure-birth process

In the spirit of the proof of theorem 1 and 2 but without strict proof, we write the increase of the average number of divisions in time interval [*t*^′^, *t*+, *dt*^′^] as

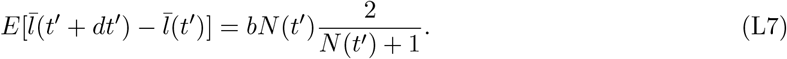

We restrict ourselves to the pure-birth process as conditioning on survival complicates the derivation. After summing over the entire time interval [0, *t*], we have

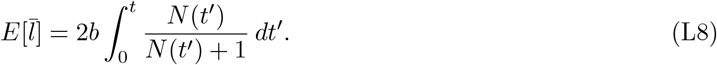

Taking *N*(*t*) = *N*_0_*e*^*bt*^, this reads

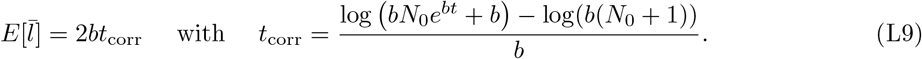

#### L.4 Comparison between mean-field solution and exact solution in the pure-birth process

##### Fixed-time solutions

Interestingly, even as time goes to infinity, there is a constant off-set between eqn. L9 and the mean field solution 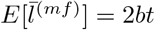 from theorem 3. We have

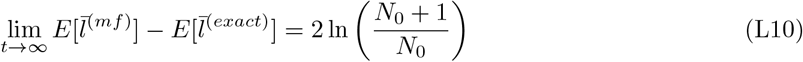

For *N*_0_ = 1, this yields a constant off-set of ln(4) ≈ 1.386, and for *N*_0_ → ∞, the difference is vanishing and we have 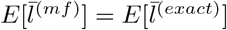.

##### Fixed-size solutions

We obtain a similar result in the fixed size condition. We transform the meanfield fixed-time solution into a fixed-size solution using according to Appenxid C such that

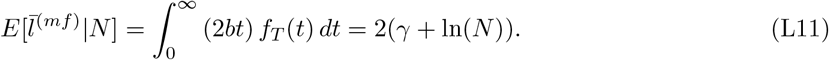

We note that *f*_*T*_ (*t*) assumes that *N*_0_ = 1. To compare this to the exact fixed-size solution, we approximate the harmonic number with its asymptotic limit 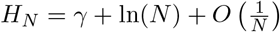 giving us

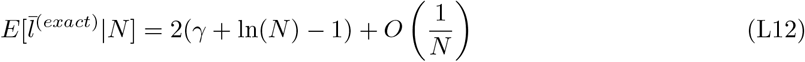

such that

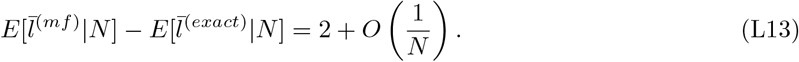

The asymptotic solution differs from the exact solution by 2, which agrees with the correction term computed for the critical branching process in ref. [36]. Further, computer simulations show that the correction term 2 is in good agreement with the birth-death process starting from a single cell.

### M Size distribution of resistant clones

We consider sensitive cells growing with rates *b*_*s*,1_ and *d*_*s*,1_ in the absence of treatment and *b*_*s*,2_ and *d*_*s*,2_ during treatment. During division of sensitive cells, each daughter cells acquires resistance mutations at rate ν ≪ 1. Resistant cells grow with rates *b*_*r*,1_ and *d*_*r*,1_ in the absence of treatment and *b*_*r*,2_ and *d*_*r*,2_ during treatment. Since the generation of resistance mutations is really small and our model does not include any other interactions between sensitive and resistant cells, we can approximate the growth of sensitive cells using growth rates *b*_*s*,1_ − *d*_*s*,1_ and *b*_*s*,2_ − *d*_*s*,2_ rather than *b*_*s*,1_ − *d*_*s*,1_ − ν and *b*_*s*,2_ − *d*_*s*,2_ − ν. We further neglect the chance that more than one resistance mutation emerge in one daughter during one division. Such events come with probability of the order ν^2^. Using these approximations, the CSD can be computed in the same way as the SFS.

#### Clone size distribution at detection

Before treatment, resistant clones are generated at rate 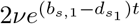 and grow to size κ with probability *p*(1 → *κ, t*_*d*_ − *t*). Following theorem 1, we have

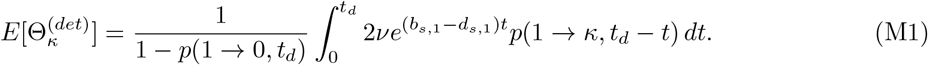

Whereas the term *p*(1 → *κ, t*_*d*_ −*t*) described the growth of resistant clones and thus must be parameterized with *b*_*r*,1_ and *d*_*r*,1_, the term 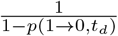 describes the conditioning on survival of the entire population that is appropriately parameterized with *b*_*s*,1_ and *d*_*s*,1_.

If sensitive and resistant cells share the same growth parameters in the absence of treatment, i.e. *b*_*s*,1_ = *b*_*r*,1_ := *b*_1_ and *d*_*s*,1_ = *d*_*r*,1_ := *d*_1_, then the CSD takes the same form as the SFS from eqn. (8), namely

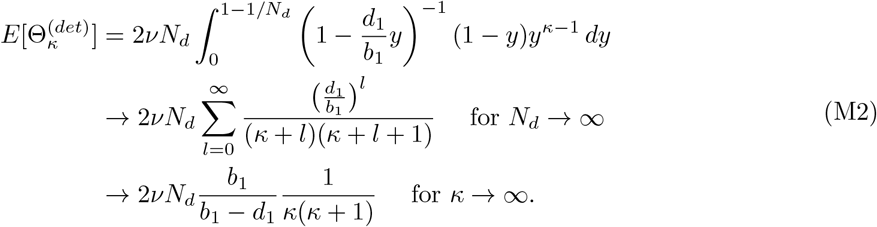

#### Clone size distribution after treatment

The CSD after treatment for time *t*_*f*_ is also obtained by adapting theorem 1. For newly emerging mutations, the relevant mutation generation comes from sensitive cells. Starting from 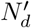 cells, the mutation generation is described by 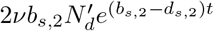. We drop the term 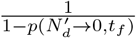 for conditioning on survival such that

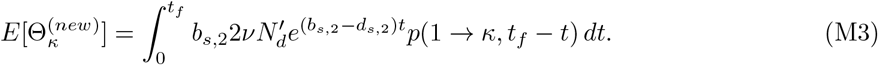

The term *p*(1 → *κ, t*_*f*_ − *t*) described the growth of resistant clones and thus must be parameterized with *b*_*r*,2_ and *d*_*r*,2_. For the CSD of clones preexisting to treatment, we again drop the conditioning on survival and obtain

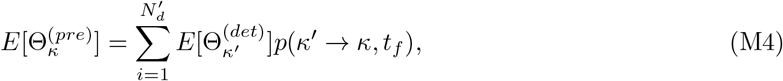

with *p*(*κ*^′^ → *κ, t*_*f*_) parameterized with *b*_*r*,2_ and *d*_*r*,2_.

To justify dropping the conditioning on survival, we note that we start from an initially large number of sensitive and resistant cells. First, it is highly unlikely that resistant cells go extinct. Second, the relevant mutational input for the CSD comes from the sensitive cell population. For small times *t*, the average growth of sensitive cells with conditioning on survival is 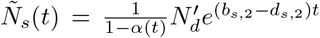 that is well approximated by the average growth of sensitive cells without conditioning 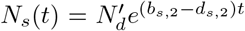. For larger *t*, when the sensitive cell population approaches the extinction threshold, there will be so few sensitive cells that the generation of new clones will be negligible, 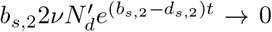. Together, this allows us to drop the term 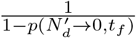 in theorem 1 while using *N*_*s*_ (*t*) rather than 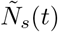 for the generation of new mutations.

### N Resistant subpopulation versus largest resistant clone

In growing populations, it is apparent that the random timing of mutations plays an important role in determining the mutant population size. Here, we ask whether the largest resistant clone dominates the total resistant subpopulation. First, we compute the arrival time of the first and second mutant and note that the second arriving mutant may outgrow the first due to genetic drift. Secondly, we derive the probability distributions for the size of the largest clone and compare this to the probability distribution for the total number of mutant cells.

We focus on clone sizes at detection since a resistant clone dominating at detection will also dominate after treatment. Therefore, we consider a birth-death process with birth rate b_1_ an death rate *d*_1_ growing from a single cell up to size *N*_*d*_, while each daughter cell acquires resistance mutation at rate ν ≪ 1 during a division. The mutation rate per division is then 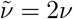.

#### N.1 Arrival time of resistant clones

In order to condition on survival of genetic drift, we can adapt the mutation rate 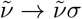, where 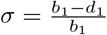 is the probability to survive drift in the linear birth-death process [15].

Considering deterministic growth of the entire population with random mutations, the number of acquired resistance mutations follows an in-homogeneous Poisson process such that the resistance mutations acquired in the time interval [τ_1_, τ_2_] is given by

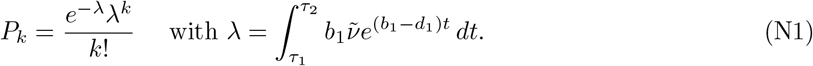

To compute the arrival time of the first mutant, we set τ_1_ = 0 and τ_2_ = *t*_1_. The probability that no mutations were acquired up to time *t*_1_ is *P*_0_. We identify 1 − *P*_0_ as the cumulative density function for the arrival time of the first surviving mutant. Taking the derivative of the cumulative density function, we obtain the probability density function for *T*_1_ that reads

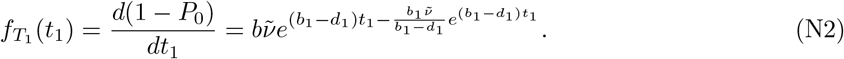

Similar results were previously reported by making approximations on the fully stochastic model (e.g. chapter 5 in ref. [15]).

Given that the first resistant clone arose at *T*_1_ = *t*_1_, the probability that no second mutations occurs until time *t*_2_ is *P*_0_ but with τ_1_ = *t*_1_ and τ_2_ = *t*_2_. Analogously to *T*_1_, we compute the probability density function for *T*_2_,

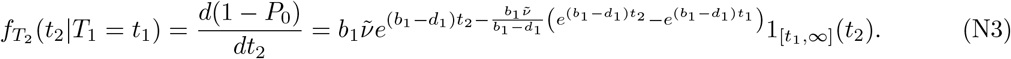

The unconditional density of *T*_2_ is obtained by marginalizing out *T*_1_,

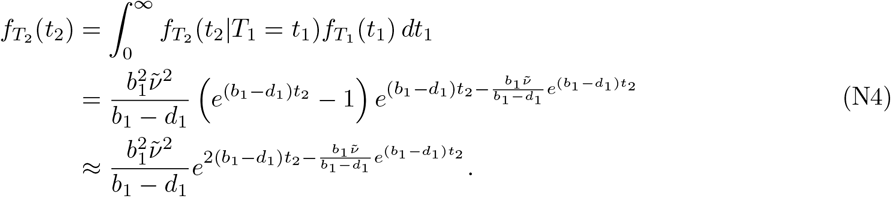

##### Inverse cumulative density function

For fig. 5A, we generate random numbers for *T*_1_ and *T*_2_. Therefore, we use inverse transformation sampling, which requires the inverse distribution function. For *T*_1_, we have

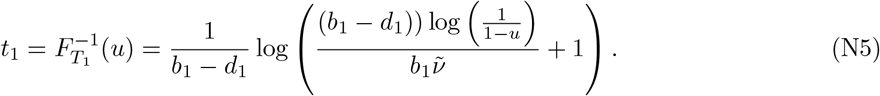

For *T*_2_ conditioned on *T*_1_ = *t*_1_, we have

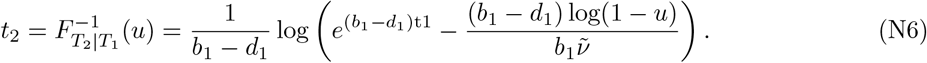

#### N.2 Size of the resistant subpopulation

In the limit 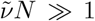, the probability to have *R* resistant cells is described by a Landau distribution [41, 42] of the form

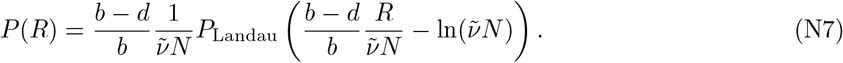

For 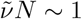, the solution is slightly more complicated but also given in ref. [42]. The fixed-time solution was computed in ref. [59] and some asymptotic solutions including selection are derived in ref. [60]. However, we focus on the case 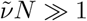.

#### N.3 Size of the largest clone

We denote the number of clones with size *κ* by Θ_*κ*_ such that 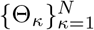 is the clone size distribution (CSD) similar to the definition of the SFS in the main text. As explained in the main text, the CSD is described by the same expression as the SFS. Since we are interested in the largest clone only that is unlikely to go extinct (should the population continue growing indefinitely), it is sufficient to consider skeleton spectrum (i.e. clones that will never go extinct) in which case the CSD takes the form described in Proposition 1 of ref. [16], namely

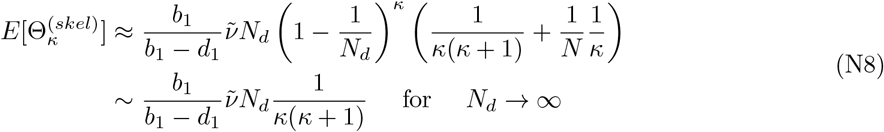

For simplicity, we take *κ* ∈ [1, ∞) on a continuous scale and approximate 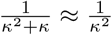. Then, the skeleton CSD can written as product of an amplitude A and probability density *f*(*κ*),

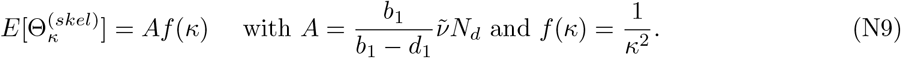

The amplitude is interpreted as the number of mutations with infinite lineage. If we take this number as fixed, then the largest clone size can be understood as order statistic. In other words, the largest clone size 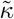 is the largest number of A clone sizes that are sampled from *f*(*κ*).

Assuming independence between clone sizes, the cumulative density function for 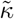 is simply the A-th power of the cumulative density function for *κ* [40]. The cumulative density function for *κ* reads 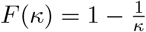 such that

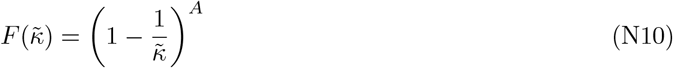

and the probability density is

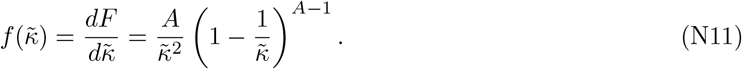

### O Site frequency spectrum upon emergence of resistance

Following the main text, we separate between mutations that occur in sensitive cells and mutations that occur in resistant cells. The sum of both contributions will make the total SFS,

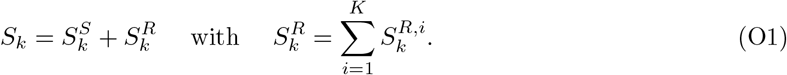

#### O.1 Mutations emerging in resistant cells

We start by analysing 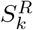, which is the sum over all mutations arising in resistant clones *i* = 1, …, *K*.

Given expected CSD *E*[Θ_*κ*_], we have

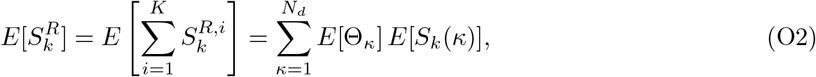

where *E*[*S*_*k*_(*κ*)] denotes the expected SFS of a resistant clone with size κ excluding mutations that were acquired prior to obtaining the resistance mutation.

Before treatment, clones grow with constant birth and death rate analogously to a homogeneous population at detection. Thus, the fixed-time SFS of a single clone with size *κ* is described by

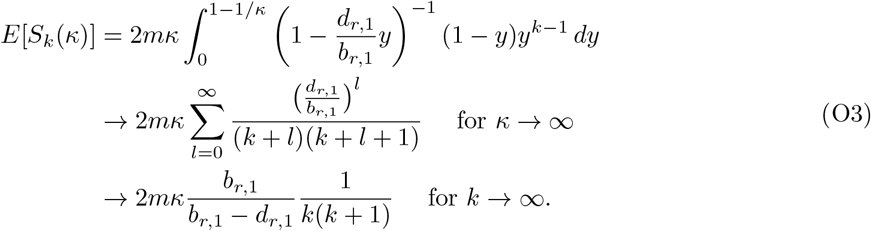

During treatment, the growth rates of the resistant clones will change and *E*[*S*_*k*_(*κ*)] is more appropriately described by homogeneous population with continued increase given by eqn. (F7) and (F8). In the case of full resistance such that *b*_*r*_ := *b*_*r*,1_ = *b*_*r*,2_ and *d*_*r*_ := *d*_*r*,1_ = *d*_*r*,2_, we can keep the expression in eqn. (O3) for the treatment phase such that

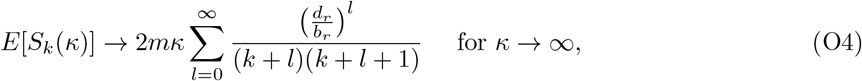

which once inserted into eqn. (O2) gives us the SFS of mutations that emerged in resistant cells. In particular, we then have

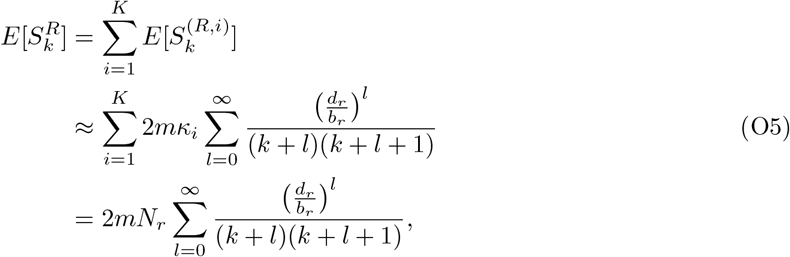

where *N*_*r*_ is the number of resistant cells. If treatment is applied long enough, there will be no more sensitive cells and we have the total population given by the resistant cells *N*_*f*_ = *N*_*r*_ giving us the neutral tail known from homogeneous populations.

#### O.2 Mutations emerging in sensitive cells

Obtaining the SFS of mutations that emerged in sensitive cells is more complicated. Neglecting the effect of resistance mutations on the growth of sensitive cells, the SFS of the sensitive cell population can be computed analogous to the homogeneous decreasing cell population whose expected SFS is described by eqn. (F1) and (F3). However, there are also mutations that emerged in sensitive cells, and are inherited to resistant cells.

For simplicity, we restrict ourselves on case that all sensitive cells went extinct. Then, mutations that emerged in sensitive cells must be clonal in at least one resistant clone, and we can write

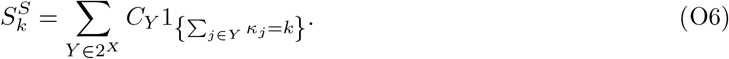

Here, the first sum goes over all possible combinations Y in the set of clone indices *X* = 1, 2, …*K* that is the power set 2^*X*^, and *C*_*Y*_ is the number of mutations shared between the selected clones.

Using theorem 3, we can connect the number of clonal mutations in a resistant clone *i*^′^ to its arrival time *t*_arrival_ ∈ [0, *t*_*d*_ + *t*_*f*_]. We have

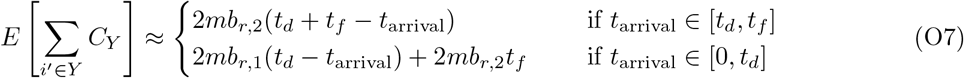

To split this predictions into individual peaks *C*_*Y*_ in the SFS, one must study the relatedness between clone *i*^′^ to other clones.

## Supplementary figures

**Figure S1:**
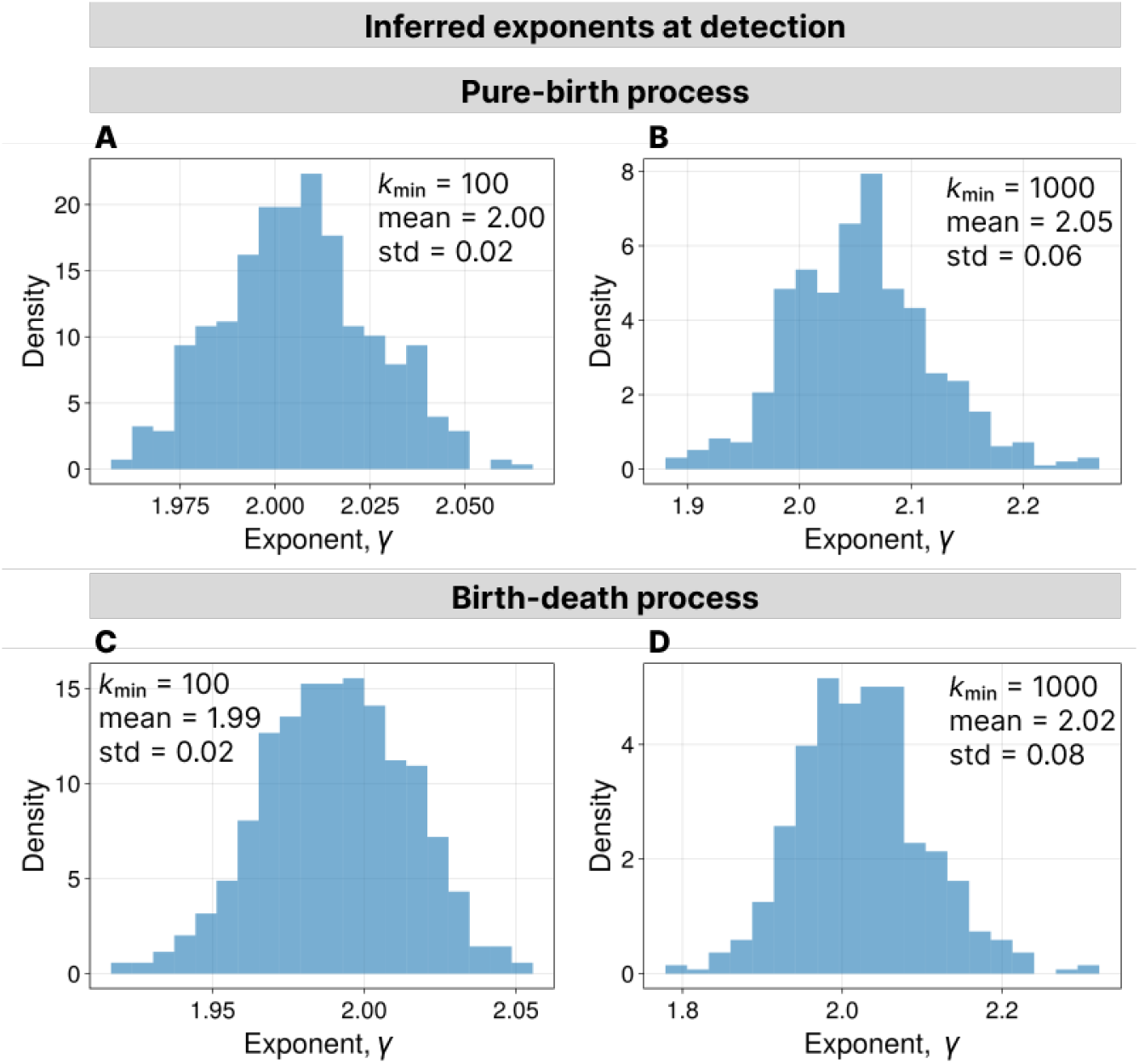
Fitted exponents to the SFS of simulated populations at detection using eqn. (37). **A**. Parameters: *b*_1_ = 1.0, *d*_1_ = 1.0, m = 2.0, *N*_*d*_ = 10^5^ and *k*_min_ = 100. **B**. Parameters: *b*_1_ = 1.0, *d*_1_ = 1.0, *m* = 2.0, *N*_*d*_ = 10^5^ and *k*_min_ = 1000. **C**. Parameters: *b*_1_ = 1.0, *d*_1_ = 0.7, *m* = 2.0, *N*_*d*_ = 10^5^ and *k*_min_ = 100. **D**. Parameters: *b*_1_ = 1.0, *d*_1_ = 0.7, *m* = 2.0, *N*_*d*_ = 10^5^ and *k*_min_ = 1000. Simulations were repeated 500 times.

**Figure S2:**
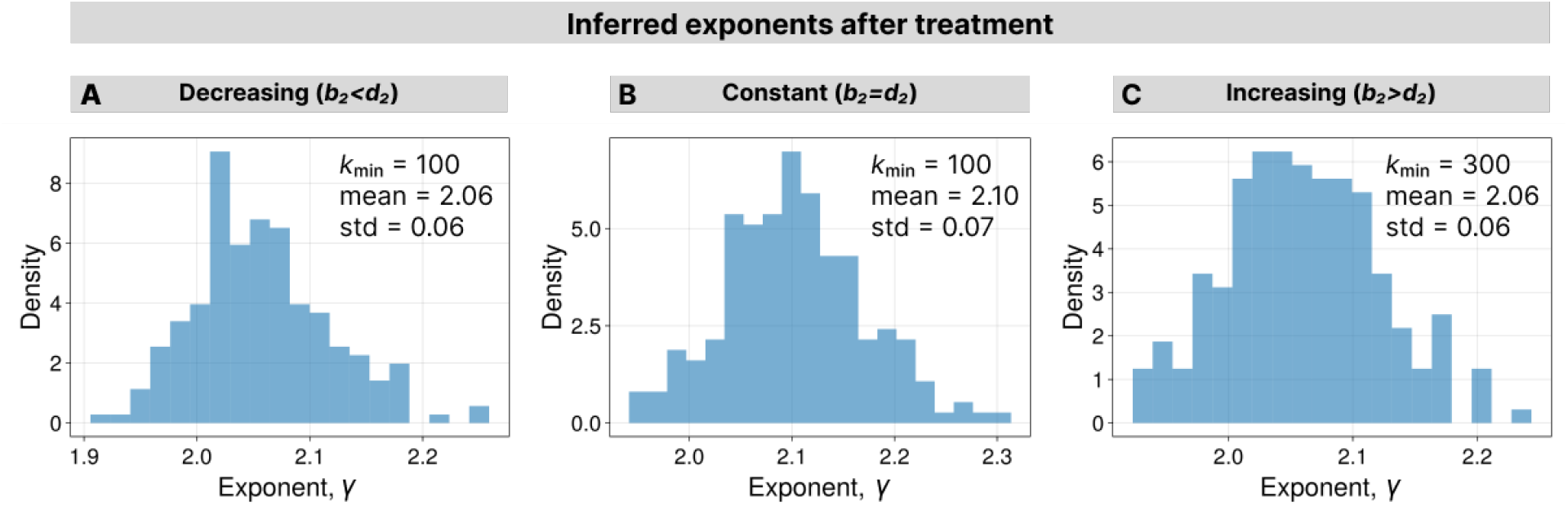
Fitted exponents to the SFS in of simulated homogeneous populations after treatment using eqn. (37). **A**. For decreasing population. Parameters before treatment: *b*_1_ = 1.0, *d*_1_ = 0.0, *m* = 2.0 and *N*_*d*_ = 10^5^. Parameters during treatment: *b*_2_ = 1.0, *d*_2_ = 3.0, *m* = 2.0 and *N*_*f*_ = 10^4^. Parameter for inference: *k*_min_ = 100. **B**. For constant population. Parameters before treatment: *b*_1_ = 1.0, *d*_1_ = 0.0, *m* = 2.0 and *N*_*d*_ = 10^5^. Parameters during treatment: *b*_2_ = *d*_2_ = 1.0, *m* = 2.0 and *t*_*f*_ = 8.0. Parameter for inference: *k*_min_ = 100. **C**. For increasing population. Parameters before treatment: *b*_1_ = 1.0, *d*_1_ = 0.0, *m* = 2.0 and *N*_*d*_ = 10^4^. Parameters during treatment: *b*_2_ = 1.0, *d*_2_ = 0.6, *m* = 2.0 and *N*_*f*_ = 3.0 × 10^4^. Simulations were repeated 200 times for each treatment scenario. Parameter for inference: *k*_min_ = 300. Simulations were repeated 200 times for each treatment scenario.

**Figure S3:**
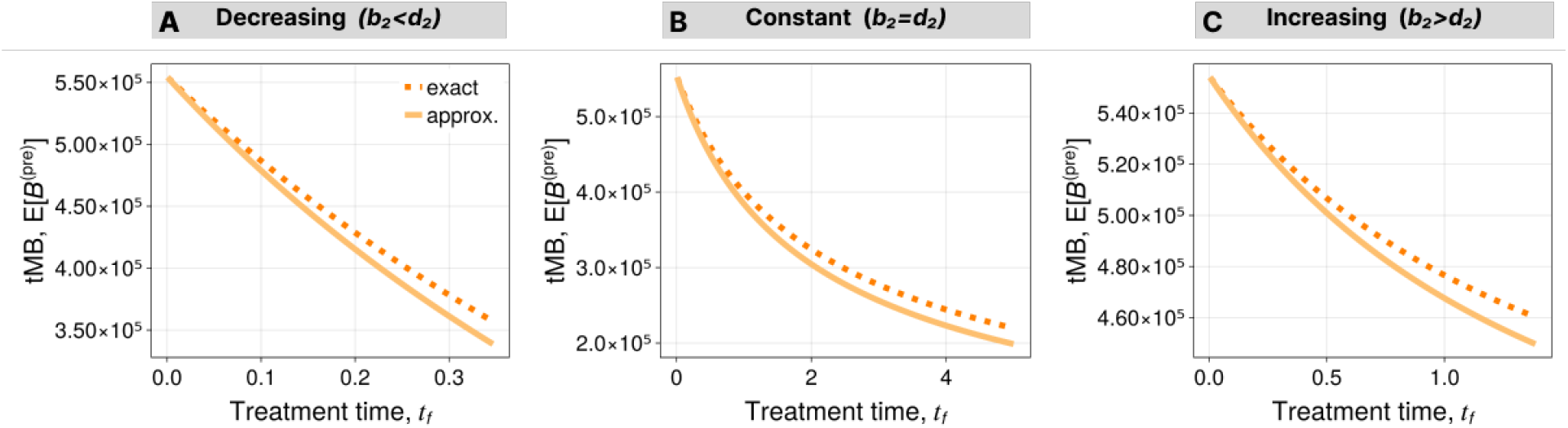
Comparison of approximate and exact expressions for the expected tMB of preexisting mutations after homogeneous treatment response. Exact tMB is given by eqn. (J2) and approximate tMB is given by eqn. (20). Parameters before treatment: *b*_1_ = 1.0, *d*_1_ = 0.5, *m* = 2.0 and *N*_*d*_ = 10^5^. Parameters during treatment for decreasing population: *b*_2_ = 1.0, *d*_2_ = 3.0, *m* = 2.0 and *t*_*f*_ varied. Parameters during treatment for constant population: *b*_2_ = *d*_2_ = 1.0, *m* = 2.0 and *t*_*f*_ varied. Parameters during treatment for increasing population: *b*_2_ = 1.0, *d*_2_ = 0.5, *m* = 2.0 and *t*_*f*_ varied.

**Figure S4:**
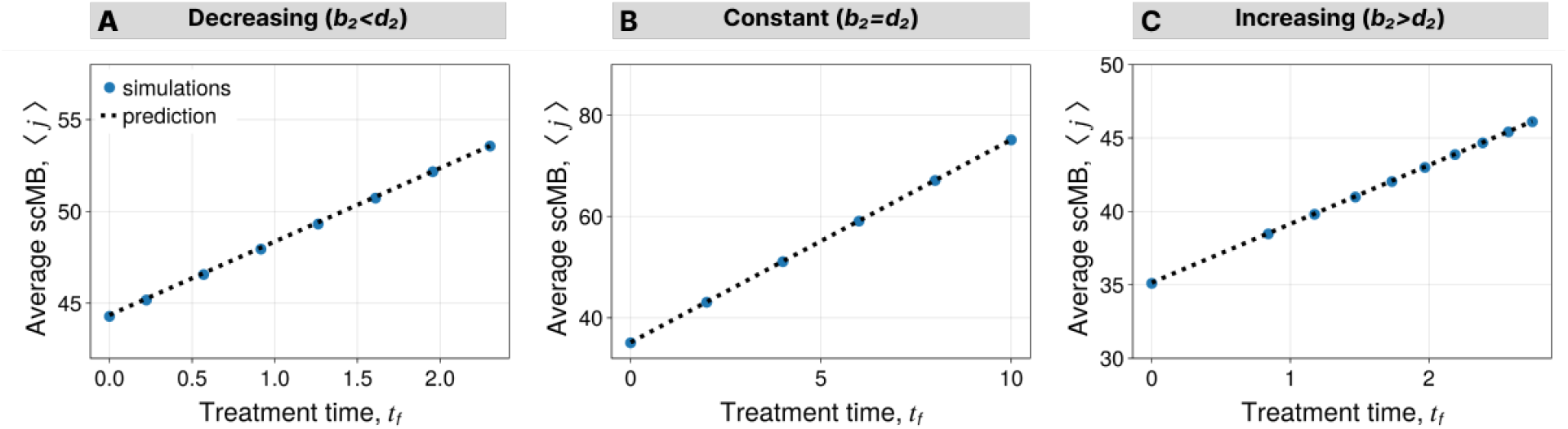
Average scMB after treatment for homogeneous population. **A**. For decreasing population. Parameters before treatment: *b*_1_ = 1.0, *d*_1_ = 0.0, *m* = 2.0 and *N*_*d*_ = 10^5^. Parameters during treatment: b_2_ = 1.0, *d*_2_ = 3.0, *m* = 2.0 and *N*_*f*_ is varied. **B**. For constant population. Parameters before treatment: *b*_1_ = 1.0, *d*_1_ = 0.0, *m* = 2.0 and *N*_*d*_ = 10^5^. Parameters during treatment: *b*_2_ = *d*_2_ = 1.0, *m* = 2.0 and *t*_*f*_ is varied. **C**. For increasing population. Parameters before treatment: *b*_1_ = 1.0, *d*_1_ = 0.0, *m* = 2.0 and *N*_*d*_ = 10^4^. Parameters during treatment: b_2_ = 1.0, d_2_ = 0.6, *m* = 2.0 and *N*_*f*_ is varied. Simulations were repeated 200 times.

**Figure S5:**
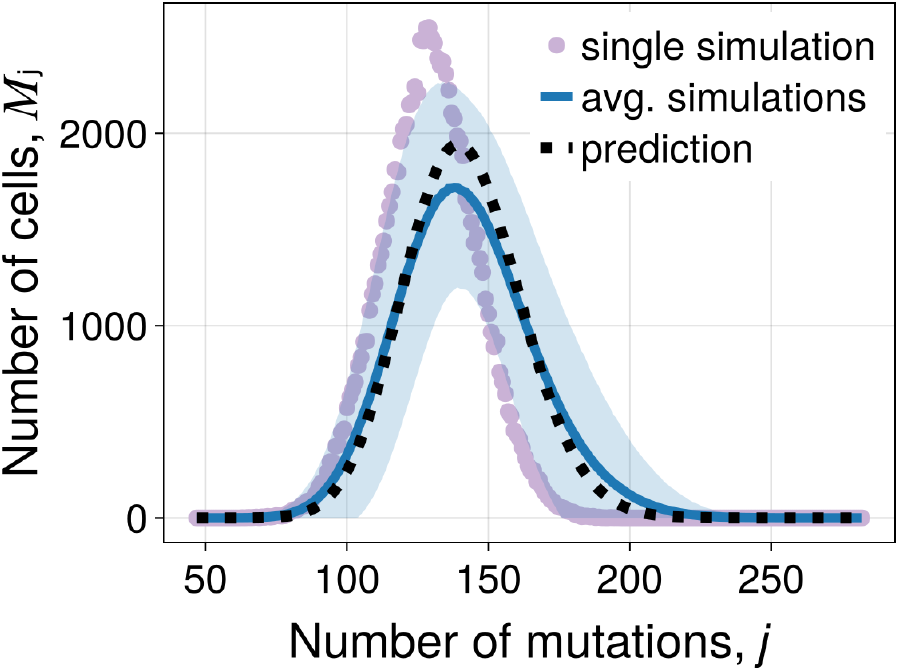
Single cell mutational burden distribution at detection for birth-death process. Parameters: *b*_1_ = 1.0, *d*_1_ = 0.7, *m* = 2.0 and *N*_*d*_ = 10^5^. Simulations were repeated 500 times.

**Figure S6:**
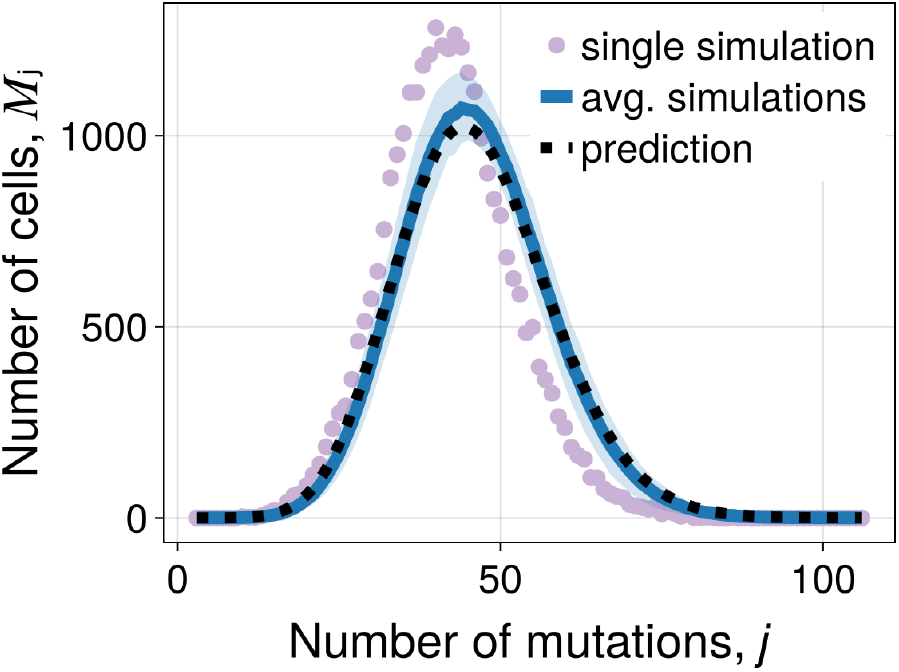
Single cell mutational burden distribution for continued increasing population. Parameters before treatment: *b*_1_ = 1.0, *d*_1_ = 0.0, m = 2.0 and *N*_*d*_ = 10^4^. Parameters during treatment: *b*_2_ = 1.0, *d*_2_ = 0.6, *m* = 2.0 and *N*_*f*_ = 3.0 × 10^4^. Simulations were repeated 200 times.

**Figure S7:**
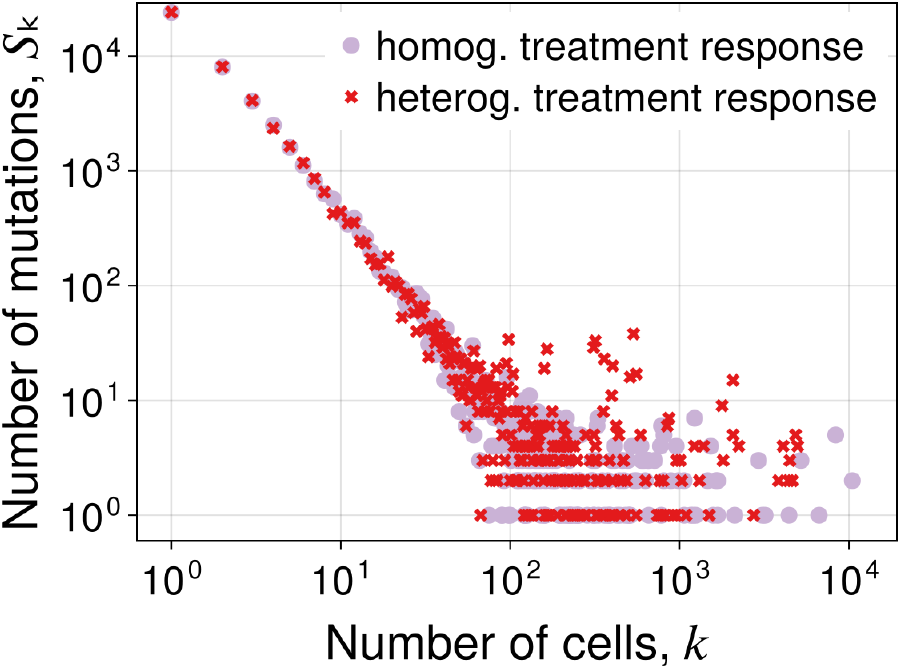
SFS of homogeneous vs heterogeneous population. For homogeneous population, we use parameters *b*_1_ = 1.0, *d*_1_ = 0.0, *m* = 2.0, *N*_*d*_ = 1.2 × 10^4^. For the heterogeneous population, we use parameters before treatment 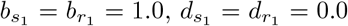, *m* = 2.0, ν = 10^−3^, *N*_*d*_ = 10 and during treatment 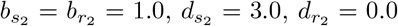, *m* = 2.0, ν = 10^−3^, *N*_*f*_ = 1.2 × 10. The shown SFS is the result of a single realization at the end of each scenario.

## Supplementary tables

**Table S1:** Parameters for fig. 2.

| Figure | $b_1$ | $d_1$ | $N_d$ | $b_2$ | $d_2$ | $N_f$ | $t_f$ | $m$ | realizations |
| --- | --- | --- | --- | --- | --- | --- | --- | --- | --- |
| 2A | 1.0 | 0.7 | $10^5$ | — | — | — | — | 2.0 | 500 |
| 2B | 1.0 | 0.0 | $10^5$ | — | — | — | — | 2.0 | 500 |
| 2C | 1.0 | 0.0 | $10^5$ | — | — | — | — | 2.0 | 500 |
| 2D | 1.0 | 0.0 | $10^5$ | 1.0 | 3.0 | $10^4$ | — | 2.0 | 200 |
| 2E | 1.0 | 0.0 | $10^5$ | 1.0 | 1.0 | — | 8.0 | 2.0 | 200 |
| 2F | 1.0 | 0.0 | $10^4$ | 1.0 | 0.6 | $3 \times 10^4$ | — | 2.0 | 200 |

**Table S2:**
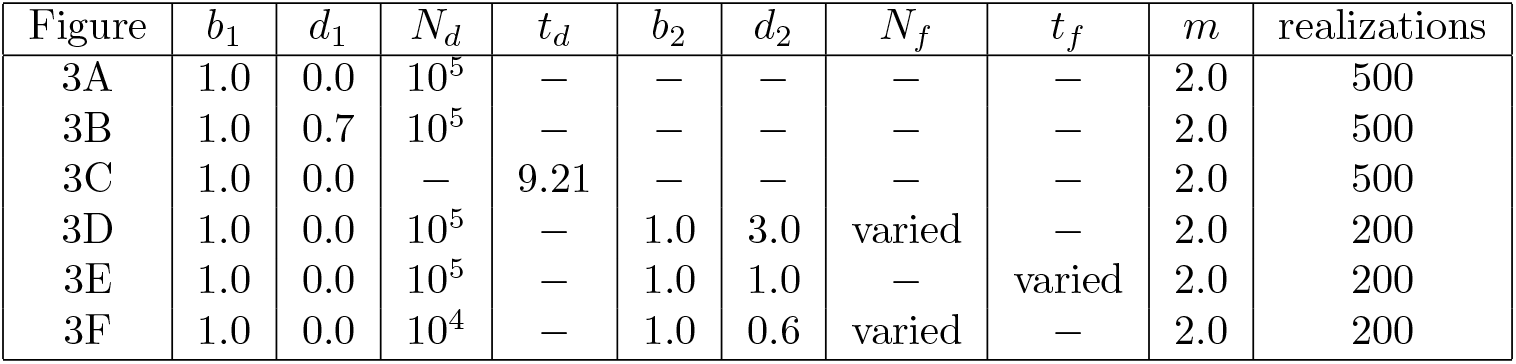
Parameters for fig. 3.

**Table S3:** Parameters for fig. 4.

| Figure | $b_1$ | $d_1$ | $N_d$ | $b_2$ | $d_2$ | $N_f$ | $t_f$ | $m$ | realizations |
| --- | --- | --- | --- | --- | --- | --- | --- | --- | --- |
| 4A | 1.0 | 0.0 | $10^5$ | — | — | — | — | 2.0 | 500 |
| 4B | 1.0 | 0.0 | $10^5$ | 1.0 | 3.0 | $10^4$ | — | 2.0 | 200 |
| 4C | 1.0 | 0.0 | $10^5$ | 1.0 | 1.0 | — | 8.0 | 2.0 | 200 |

**Table S4:** Parameters for fig. 5. We set *b*_*s*,1_ = *b*_*r*,1_ = *b*_1_ and *d*_*s*,1_ = *d*_*r*,1_ = *d*_1_.

| Figure | $b_1$ | $d_1$ | $N_d$ | $t_d$ | $b_{s,2}$ | $d_{s,2}$ | $b_{r,2}$ | $d_{r,2}$ | $N_f$ | $m$ | $\nu$ | realizations |
| --- | --- | --- | --- | --- | --- | --- | --- | --- | --- | --- | --- | --- |
| 5A | 0.14 | 0.13 | — | 2555 | — | — | — | — | — | — | $10^{-7}$ | $10^6$ |
| 5B | 1.0 | 0.5 | $10^5$ | — | — | — | — | — | — | — | $10^{-3}$ | 500 |
| 5C | 0.14 | 0.13 | varied | — | — | — | — | — | — | — | $10^{-7}$ | 500 |
| 5D | 1.0 | 0.0 | $10^4$ | — | 1.0 | 3.0 | 1.0 | 0.0 | $2.0 \times 10^4$ | 2.0 | $10^{-3}$ | 200 |
| 5E | 1.0 | 0.0 | $10^4$ | — | 1.0 | 3.0 | 1.0 | 0.0 | $2.0 \times 10^4$ | 2.0 | $10^{-3}$ | 200 |
| 5F | 1.0 | 0.0 | $10^4$ | — | 1.0 | 1.5 | 0.7 | 0.0 | $1.2 \times 10^4$ | 2.0 | $10^{-3}$ | 1 |

## Notes

### Competing Interest Statement

The authors have declared no competing interest.

